# Deciphering BRCAness Phenotype in Cancer: A Graph Convolutional Neural Network Approach with Layer-wise Relevance Propagation Analysis

**DOI:** 10.1101/2024.06.26.600328

**Authors:** Jingyu Yang, Hryhorii Chereda, Jürgen Dönitz, Annalen Bleckmann, Tim Beißbarth

## Abstract

**Background:** Cancer variability among patients underscores the need for personalized therapy based on genomic understanding. BRCAness, characterized by vulnerabilities similar to BRCA mutations, particularly in homologous recombination repair, shows potential sensitivity to DNA-damaging agents like PARP inhibitors, highlighting it’s clinical significance.

**Methods:** We employed Graph Convolutional Neural Networks (GCNNs) with Layer-wise Relevance Propagation (LRP) to analyze gene expression data from the TCGA Pan-Cancer dataset. The study compared the efficacy of GCNNs against traditional machine learning models and differential gene expression analysis, focusing on their ability to elucidate complex genomic interactions defining BRCAness.

**Results:** Differential Gene Expression (DGE) analysis proved limited in capturing the nuances of BRCAness. In contrast, GLRP significantly identified genes related to transcription regulation and cancer processes, emphasizing the phenotype’s complexity. Gene Set Enrichment Analysis (GSEA) highlighted crucial pathways like Nuclear Receptors signaling, Cellular Senescence, and ESR-mediated signaling, underscoring their roles in BRCAness and therapeutic potential.

**Conclusion:** GLRP outperformed traditional approaches in analyzing BRCAness, providing deep insights into transcriptional and oncogenic processes critical to the BRCAness phenotype. Our findings suggest new directions for developing targeted and personalized cancer treatments, leveraging intricate molecular interactions associated with BRCAness.

## 1 Introduction

Cancer remains one of the leading causes of mortality worldwide, with its complexity and variability across patients posing significant challenges for treatment. At the heart of personalized cancer therapy is the understanding of cancer genomics, which seeks to unravel the genetic underpinnings of cancer phenotypes to guide treatment decisions. Among these phenotypes, a crucial one to understand is related to the BRCA genes. Initially, mutations in BRCA1 and BRCA2 are well-known for their role in breast cancer, where they have led to approved therapies [1]. These mutations can be either germline, inherited from a parent, or somatic, acquired during a person’s lifetime [2]. Germline BRCA1/2 mutations significantly increase the risk of developing breast, ovarian, and other cancers [3]. Somatic BRCA1/2 mutations are also prevalent in various cancer types, occurring in approximately 5–7% of ovarian cancers [3].

Building on this understanding, the concept of BRCAness has emerged. It extends beyond BRCA1 and BRCA2 mutations and describes tumors that, despite lacking mutations in these genes, exhibit similar molecular vulnerabilities [4]. This is particularly evident in the context of deficiencies in homologous recombination repair, a high-fidelity pathway responsible for repairing double-strand DNA breaks [5–7]. This phenotype suggests potential sensitivity to DNA damaging agents, such as PARP inhibitors, which exploit the synthetic lethality concept by targeting the base excision repair pathway in cells with HRR deficiencies [8, 9]. Identifying and comprehensively understanding BRCAness is crucial for the molecular tumor board, which aims to integrate genomic data, molecular characteristics, and clinical information to guide personalized treatment strategies [10]. By identifying BRCAness in tumors, even in the absence of BRCA1/2 mutations, clinicians can recommend targeted therapies, such as PARP inhibitors, to a broader range of patients, thereby potentially improving outcomes and tailoring treatment approaches [11, 12].

Previous research has progressively illuminated the complex genetic and molecular landscape defining BRCAness, identifying specific gene expressions and mutational signatures associated with this phenotype across various cancer types. Notably, Chen’s study underscores the role of CXCL1 and LY9 upregulation in ovarian cancer BRCAness [13], whereas Goundiam et al. [14] observed CCNE1 and BRD4 overexpression in non-BRCAness high-grade ovarian carcinoma. Konstantinopoulos’s findings [15] further establish a link between BRCAness gene expression profiles and enhanced responses to platinum and PARP inhibitors, underscoring the therapeutic relevance of these molecular markers. Additionally, Telli’s research [16] on triple-negative breast cancers emphasizes the predictive value of the Homologous Recombination Deficiency (HRD) score for platinum-based therapy responsiveness, a sentiment echoed by Murai’s [5] discussion on HRD score thresholds in selecting HR-deficient tumors or BRCAness. How’s examination [17] of HRD score alternatives for PARPi prediction adds another layer to the multifaceted approach required in assessing BRCAness. Davies’ implementation of HRDetect [18], which uses mutational signatures to identify BRCA1/2 deficiencies, proposes an expanded cohort of breast cancer patients who could benefit from PARP inhibitors. Despite these advances, the field still grapples with a lack of comprehensive analysis across cancer types, highlighting an urgent need for integrated studies that can consolidate these findings into a clearer understanding of BRCAness, thereby guiding more effective therapeutic interventions.

Recent advancements in computational biology and machine learning have opened new avenues for deciphering complex cancer phenotypes like BRCAness or cancer subtypes [19]. Graph Convolutional Neural Networks (GCNNs) have emerged as a powerful tool for analyzing graph-structured genomic data [20], enabling the integration of gene expression profiles with functional interaction networks [21]. By leveraging the topological information inherent in these networks, GCNNs can uncover intricate patterns and dependencies that traditional machine learning approaches may overlook [22]. Furthermore, the application of Graph Layer-wise Relevance Propagation (GLRP) in conjunction with GCNNs offers an interpretable framework for identifying the most contributory genes driving the BRCAness phenotype. This study aims to harness the power of GCNNs and GLRP to conduct a comprehensive, pan-cancer analysis of the BRCAness phenotype. By integrating multi-dimensional genomic data from the TCGA Pan-Cancer dataset, including gene expression profiles, Homologous Recombination Deficiency (HRD) scores [16], and somatic mutation information, we seek to unravel the complex genetic underpinnings of BRCAness across diverse cancer types. Through a rigorous labeling strategy that incorporates HRD scores and pathogenic mutations in key BRCAness genes, we aim to capture the canonical definition of BRCAness for interpretative analysis.

Moreover, by comparing the performance and feature importance of GCNNs with traditional machine learning models and differential gene expression analysis, we aim to demonstrate the superiority of graph-based approaches in capturing the nuances of the BRCAness phenotype. Ultimately, this study seeks to advance our understanding of the genetic architecture underlying BRCAness, identifying key genes, pathways, and biological processes that contribute to this phenotype. By leveraging cutting-edge computational techniques and integrating diverse genomic data, we aim to pave the way for more targeted and personalized cancer therapies, exploiting the molecular features associated with BRCAness to improve patient outcomes and guide potential treatment decisions.

## 2 Materials and Methods

### 2.1 Model Development

The analysis workflow shown in Figure 1 for the BRCAness phenotype leverages the capabilities of Graph Convolutional Neural Networks (GCNNs) in conjunction with Layer-wise Relevance Propagation (LRP) which was depicted in Chereda’s work [22]. GCNN+LRP applies to graph-structured data and allows not only for data point-specific explanations but also for model-wide scoring and selection of features [23]. We utilized the functional interaction networks from Reactome to structure patient’s gene expression data [24]. GCNN performs classification of patients, by mapping their gene expression data, denoted as *X ∈ R^m×n^*, to a target variable *Y ∈ R^m^*, which represents the BRCAness status. Here, *m* is the number of samples (patients) and *n* is the number of features (genes). The gene expression data is structured in an undirected graph *G* = (*V, E, A*), where *V* and *E* represent the sets of vertices (genes) and edges (interactions), respectively, and *A* is the adjacency matrix. The number of vertices corresponds to the number of genes. The values in the gene expression matrix *X* are interpreted as the graph signal.

**Fig. 1.**
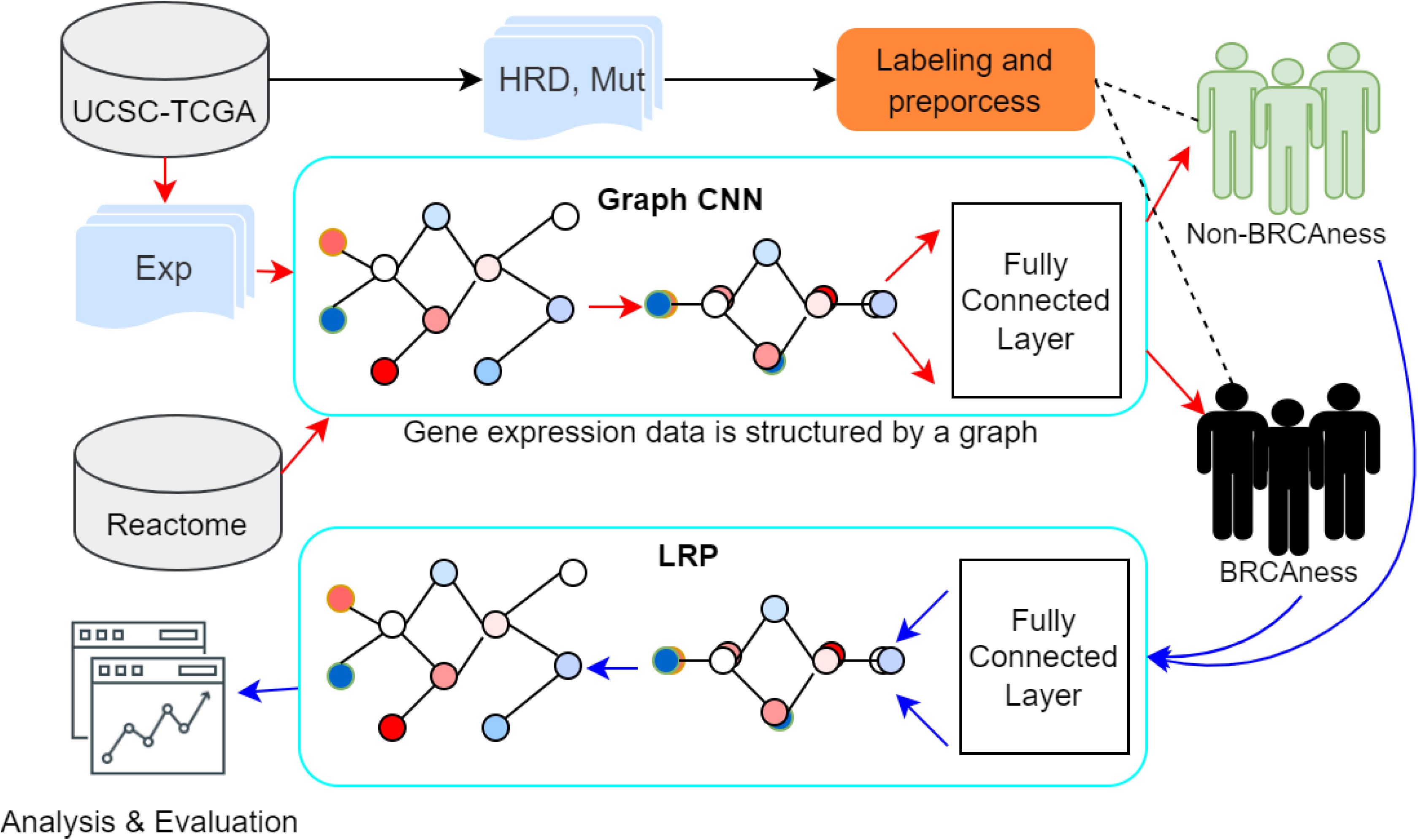
The workflow of BRCAness Phenotype analysis employs Graph Convolutional Neural Networks (GCNNs) and Layer-wise Relevance Propagation (LRP). Gene expression data, somatic mutations, and HRD (Homologous Recombination Deficiency) scores are collected from TCGA samples in the UCSC-TCGA repository. Each sample is labeled as BRCAness or non-BRCAness based on HRD scores and the mutation status of BRCAness associated genes. For each patient, gene expression data is constructed into a graph based on gene functional interactions (FI), where each node represents a gene expression profile, and this network is consistent for each patient. The gene expression values of the patient are assigned to each node of the gene functional interaction network, thus representing the patient as a graph signal. The GCNN (Graph Convolutional Neural Network) performs graph convolutions and classifies patients as BRCAness or non-BRCAness. LRP (Layer-wise Relevance Propagation) propagates relevance from the predicted labels back to the input features (nodes of the gene functional network). The top 100 features with the highest relevance are extracted for further analysis and evaluation.

GCNNs depart from traditional CNNs by employing data that resides on these graphs. The essence of GCNN lies in graph convolution, which is implemented through the graph Laplacian *L* = *D − A*, with *D* as the weighted degree matrix and *A* as the adjacency matrix. The graph convolution captures localized patterns within the graph signal and is mathematically expressed through eigendecomposition of the Laplacian *L* = *U* Λ*U ^T^*. This enables a filtering operation, where *x, y ∈ R^m^* are graph signals, and the filter *h_θ_*(Λ) is a function of eigenvalues of *L*, which are treated as graph frequencies.

The spatial localization of filters is achieved through a polynomial parametrization, defined by the expression:

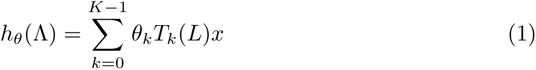

where *θ ∈ R^k^* is a vector of parameters, and *T_k_*(*x*) denotes the Chebyshev polynomial of order *k*. This enables efficient computation of the GCNN filtering operation, allowing it to scale to large graphs.

Once the GCNN model has been trained and validated using a 5-fold cross-validation scheme to ensure robust predictive performance, LRP is employed on correctly classified samples in the test set. LRP works by backpropagating relevance scores from the output layer to the input features, determining the contribution of each gene to the BRCAness phenotype. Let *i* and *j* be single neurons at two consecutive layers at which the relevance should be propagated from *j* to *i*. The LRP algorithm utilizes the z-rule for relevance propagation:

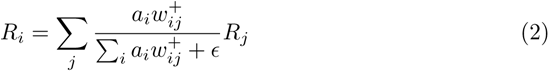

where 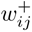 corresponds to the positive weights *w_ij_* and *ɛ* stabilizes numerical computations. This equation guides the backward redistribution of relevance from the output layer back to the input layer, attributing significance to the individual contributions of the genes. The iterative application of LRP across the network layers illuminates the decision-making process of the GCNN, allowing us to pinpoint gene features with the highest impact on the BRCAness classification outcome, thus offering valuable insights into the genomic architecture underlying this phenotype.

In contrast to traditional machine learning models such as Logistic Regression (LR), Support Vector Machine (SVM), Decision Tree, Gaussian Naive Bayes (GNB), and Random Forest ([25–28]), the application of Graph Convolutional Neural Network (GCNNs) to the BRCAness phenotype analysis offers a significant advantage by inherently incorporating the molecular network structure into the learning process. Whereas conventional methods might overlook the complex interaction between genes, GCNNs explicitly model these interactions through the graph’s edges and vertices, leveraging the adjacency matrix *A* within the graph Laplacian *L* = *D − A* to capture the localized patterns of gene expression. This structural integration allows GCNNs to not only consider the individual gene features but also their contextual significance in the gene expression network. Consequently, GCNNs provide a more nuanced understanding of the BRCAness status, as they are able to discern the intricate biological pathways that traditional machine learning models, which treat features independently, cannot inherently capture.

### 2.2 Data Source

The TCGA Pan-Cancer (PANCAN) genomic information was collected from the resource UCSC Xena [29], covering 33 cancer types. This repository represents an extensive collection of multi-dimensional cancer genomics data, gathered from a large cohort of patients across various cancer types. For our analysis, we extracted three critical types of data: gene expression profiles, Homologous Recombination Deficiency (HRD) scores, and somatic mutation information. The gene expression data provides a comprehensive landscape of gene activity across different tumors, offering insights into diverse molecular mechanisms at play. HRD scores, quantifying deficiencies in homologous recombination repair mechanisms, are pivotal for identifying potential BRCAness characteristics in tumors. Lastly, the somatic mutation data sheds light on the individual genetic alterations driving cancer progression.

Gene functional interaction derived from the Reactome [24] database was used for constructing the structure of gene expression data. Reactome is renowned for its comprehensive and accurately curated data on human biological processes, including molecular interactions. We leveraged its wealth of functional interaction information to construct a network that linked genes through their functional relationships, rather than just expression similarities. This network, which incorporated both direct and inferred gene interactions, was a key element in constructing our graph convolutional neural network (GCNN).

Additionally, genes associated with transcription regulation were collected using the Gene Ontology term GO:0006355 [30, 31]. Cancer-related genes were curated from the Cancer Gene Census, part of the COSMIC database [32], which is renowned for its comprehensive catalog of genes with established links to cancer. For genes associated with BRCAness, we referred to the comprehensive review by Guo et al. [6], which provides an in-depth exploration of genes implicated in homologous recombination deficiencies.

### 2.3 Sample Labelling and Preprocessing

#### 2.3.1 Strategy for BRCAness and Non-BRCAness Labelling

The labeling strategy for BRCAness and Non-BRCAness is grounded in the foundational concept of BRCAness, which refers to a tumor’s defect in the homologous recombination repair (HRR) pathway, often resembling the loss of BRCA1 or BRCA2 function [5]. Given the complexity and multifaceted nature of BRCAness, encompassing various dimensions of tumor biology, a single definitive “gold standard” for its identification remains elusive. Consequently, we have devised a robust labeling strategy, drawing upon existing research on BRCAness [5, 33]. This approach integrates the Homologous Recombination Deficiency (HRD) score—an aggregate of three genomic instability measures: HRD-LOH (loss of heterozygosity), HRD-TAI (telomeric allelic imbalance), and HRD-LST (large-scale state transitions)—with the mutation status of key BRCAness genes. Prior studies, such as those by Chai [34], Li [35], and Telli [16], indicate that HRD scores of 33 and 42 are commonly used thresholds for defining HR deficiency, with 42 representing a more stringent criterion and 33 a less restrictive one. Furthermore, pathogenetic mutation information associated with BRCAness genes, derived from Guo’s review [6], includes genes such as ATM, MYC, TP53, BAP1, BARD1, BLM, BRCA1, BRCA2, and others.

The patient labeling process for BRCAness/Non-BRCAness, as illustrated in Figure 2, unfolds as follows:

1. HRD score, and pathogenetic mutations linked to BRCAness of each patient are collected from the TCGA-PAN dataset and Guo’s supplementary data ([6]). Patients lacking any of these data are excluded.
2. In the absence of BRCA1 or BRCA2 mutations, those with an HRD score of 42 or higher are classified as BRCAness.
3. In the absence of BRCA1 or BRCA2 mutations, those with an HRD score of 42 or higher are classified as BRCAness.
4. For cases not meeting the above criteria, patients with an HRD score of 33 or above, coupled with pathogenic mutations in key BRCAness genes (excluding BRCA1 or BRCA2), are also labeled as BRCAness.
5. Patients who do not fulfill any of the aforementioned criteria are designated as Non-BRCAness.

**Fig. 2.**
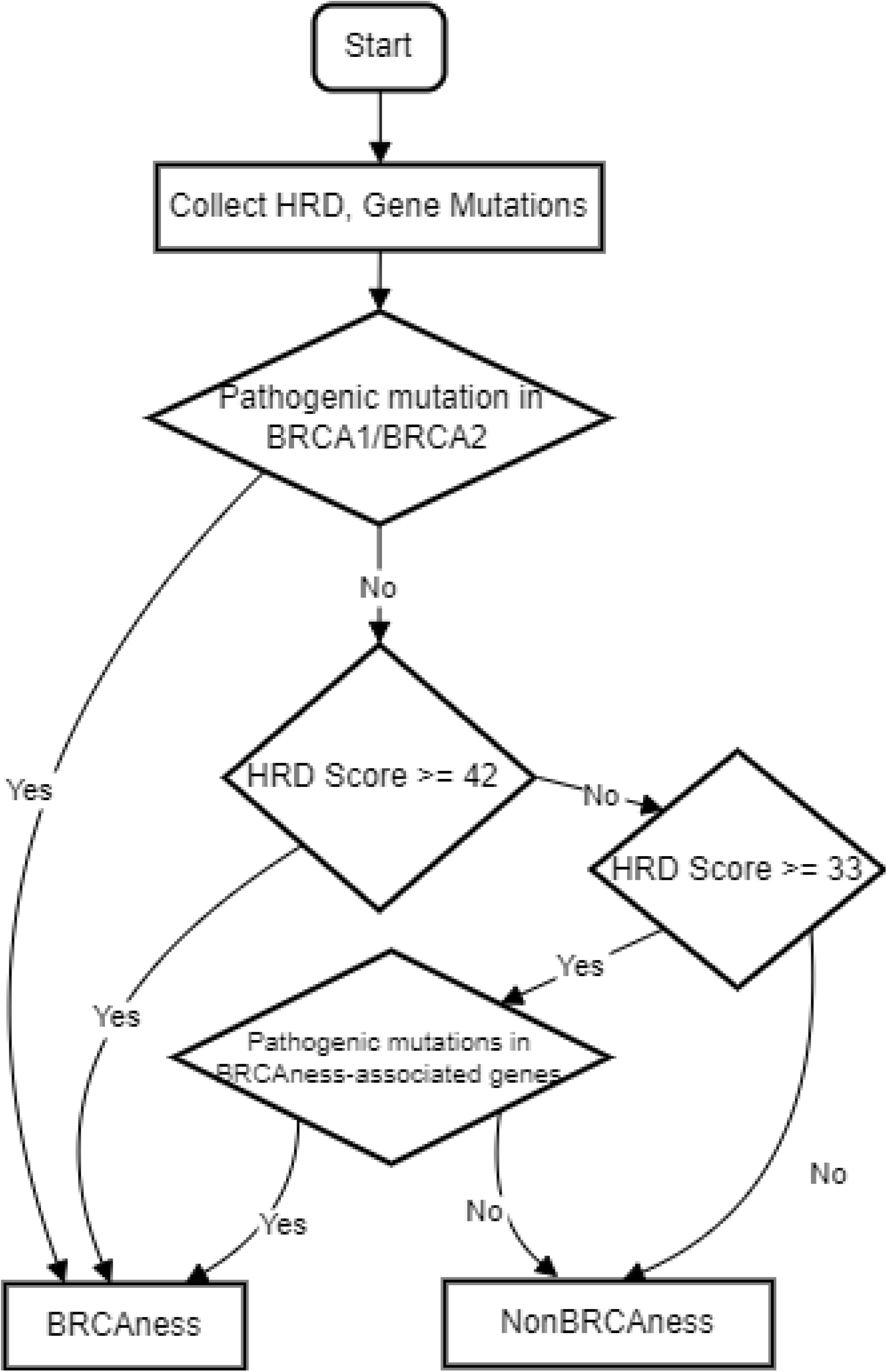
The workflow of BRCAness Labeling strategy.

Through this hierarchical and conditional strategy, the labeling captures not only the canonical definition of BRCAness, reflecting a defect in the HRR pathway similar to BRCA1 or BRCA2 loss but also a broader concept of HR deficiency.

#### 2.3.2 Data Balancing and Cancer Type Selection

In the labeling preprocessing stage, our dataset comprised 32 cancer types, totaling 8,156 labeled patients, after accounting for the absence of Acute Myeloid Leukemia (LAML) in the BRCAness gene pathogenetic mutation files. A critical observation, as depicted in figure 3, is the extreme skewness in the original labeling of TCGA samples. In several cancer types, the prevalence of Non-BRCAness cases markedly outweighed that of BRCAness cases, a disparity that could significantly impede the performance of any machine learning methodology. To address this data imbalance, we implemented a resampling strategy focused on dataset balancing.

**Fig. 3.**
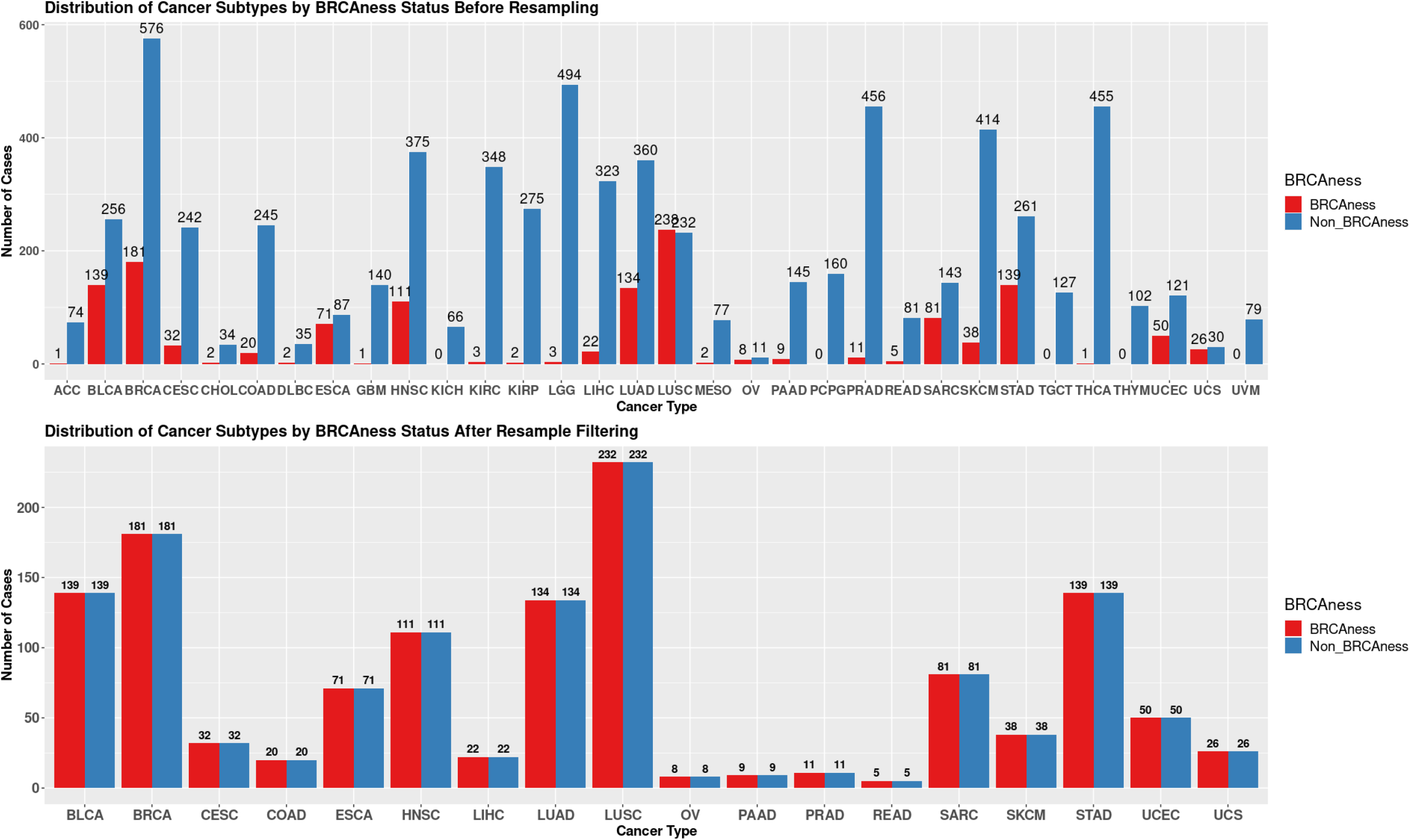
TCGA dataset with BRCAness/Non-BRCAness Labelling before and after data resampling.

Our approach involved downsampling across all cancer types, aimed at equalizing the sample sizes between the smaller groups within each type (BRCAness/Non-BRCAness). This process was critical to ensure a more balanced representation and mitigate the bias inherent in the original dataset. Subsequently, we also excluded cancer types with insufficient representation, specifically those with fewer than 5 cases in either category (BRCAness or Non-BRCAness). This exclusion criterion was essential to maintain the integrity and reliability of our analysis, as extremely low sample sizes could compromise the validity of our findings and the efficacy of our machine learning models. Upon obtaining a dataset of 2,618 resampled and labeled samples, we proceeded to divide it into training and test sets. This division was facilitated through the use of a train-test split, with stratification based on a combination of cancer type and BRCAness label. This strategy ensured that both sets accurately reflected the composition of the entire dataset. The final allocation resulted in a training set comprising 2,094 samples and a test set with 524 samples (Fig. 3).

### 2.4 Comparative Analysis of Gene Contributions to BRCAness

Following the labeling and preprocessing of samples, we applied Graph Convolutional Neural Network (GCNN) training coupled with Layer-wise Relevance Propagation (LRP) analysis. The Graph Layer-wise Relevance Propagation (GLRP) analysis yielded a results matrix detailing the relevance scores for each gene across patients in the test set. To focus our analysis, we retained only those samples that were correctly classified. Subsequently, we computed the absolute mean of the LRP scores for each gene. This process enabled us to identify and select the top 100 genes as the most contributory factors in the BRCAness phenotype, highlighting their potential significance in understanding and diagnosing this condition. To visually represent these findings, gene expression heatmaps and gene interaction networks for each cancer type were constructed.

In parallel, the resampled dataset underwent training using traditional machine learning models, including Logistic Regression, Support Vector Classifier, Decision Tree, Random Forest, and Gaussian Naive Bayes, to allow for comparative analysis. The subsequent feature importance analysis for these models was conducted as follows: For tree-based models like Random Forest and Decision Tree, gene feature importance was quantified based on the decrease in node impurity, with genes facilitating the most critical splits deemed most important. In Logistic Regression and Linear Support Vector Classification (SVC), gene feature importance was derived from the coefficients attributed to each gene, where the coefficients represent the change in log odds (for Logistic Regression) or the margin (for SVC) for a unit change in gene expression. The top 100 genes identified by each model were then selected for a comparative analysis with genes associated with BRCAness. Additionally, differential gene expression analysis [36] was applied to the resampled dataset to investigate the presence of differentially expressed genes within the context of BRCAness.

## 3 Results

### 3.1 Differential Gene Expression Analysis

The differential expression gene (DGE) analysis was conducted on the resampled and labeled dataset, employing a threshold of *P−value <* 0.05 and *fold change >* 1. This analysis was visualized in a volcano plot (Fig. 4). Intriguingly, the results revealed that none of the differentially expressed genes intersected with the genes associated with the BRCAness phenotype. This observation suggests that traditional DGE analysis may encounter challenges in uncovering the nuances of the BRCAness phenotype.

**Fig. 4.**
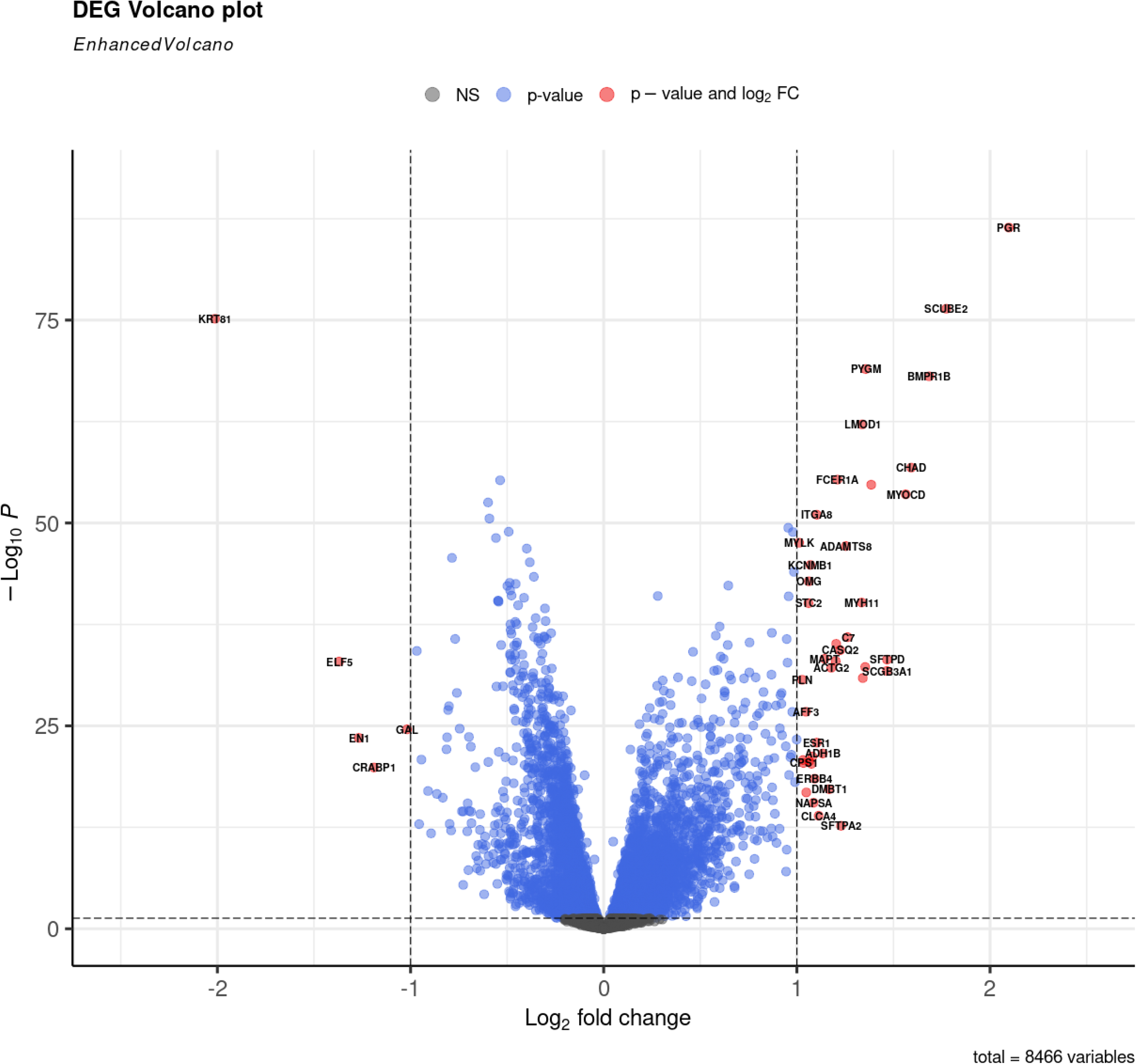
DGE volcano plot for the TCGA BRCAness/Non-BRCAness resampled dataset.

### 3.2 Performance of the GCNN Model

In the comparative analysis of machine learning models for classifying BRCAness in the scopes of 5-fold cross-validation, the Graph Convolutional Neural Network (GCNN) model emerged as the top performer with an average accuracy of 0.7427, outshining Logistic Regression, Support Vector Classifier, Decision Tree, Random Forest, and Gaussian Naive Bayes (Table 1.). It also achieved the highest precision and F1 Score, signifying its ability to more accurately identify and classify BRCAness cases. A contributing factor to the GCNN model’s success is its integration of gene interaction networks, which enables it to capture complex patterns in genomic data that traditional models may overlook.

**Table 1.**
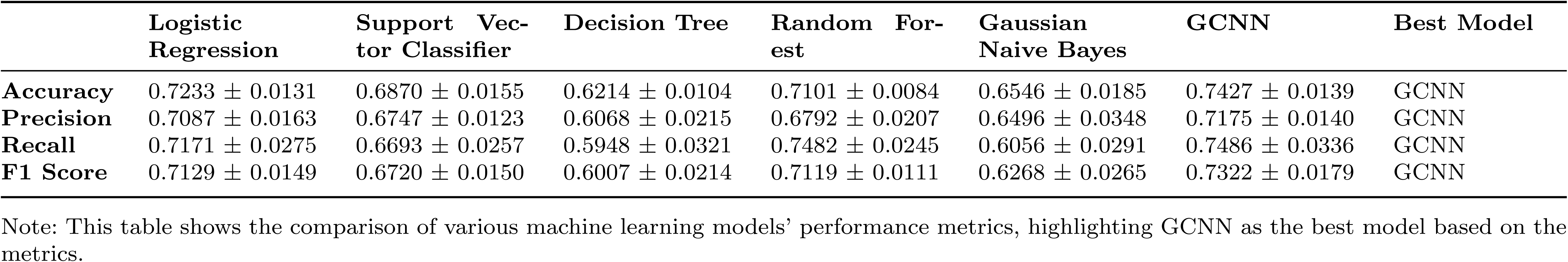
Comparison of Machine Learning Model Performance Metrics in the scopes of 5-fold cross-validation: The table summarizes the accuracy, precision, recall, and F1 score for six different models (Logistic Regression, Support Vector Classifier, Decision Tree, Random Forest, Gaussian Naive Bayes, GCNN).

### 3.3 Delineating BRCAness Through Top LRP Genes Expression and Functional Interactions

The top contributing genes were identified through Layer-wise Relevance Propagation (LRP) analysis to further investigate the BRCAness phenotype. To achieve a deeper understanding, we constructed a gene expression heatmap and a corresponding gene interaction network for cancer types with sufficient cases to hold statistical power in the test set.

The expression heatmap split by cancer type (Fig. 5) illustrates the expression level of the top 100 genes with the highest LRP scores across various cancer types, categorized by BRCAness and Non-BRCAness phenotypes. Genes are ordered based on the median difference in expression levels between BRCAness and Non-BRCAness groups, with annotations on the right indicating the median expression within each phenotype. Color gradations indicate gene expression, with red representing higher and blue indicating lower expression levels.The uppermost annotations categorize samples by cancer type, gender, age, and BRCAness status, providing a multifaceted view of the data. To the right, genes implicated in cancer, as per the Cancer Gene Census from COSMIC [32], are marked in red; those involved in gene transcription regulation, extracted from Gene Ontology (GO:0006355), are highlighted in green; and a blue annotation highlights the rank of genes based on their LRP scores. The color-coded gene symbols on the right reinforce this classification, with green for transcription regulators, red for cancer genes, purple for genes with dual roles, and black for genes that are neither transcription regulators nor identified cancer genes. The heatmap (Fig. 5) clearly demonstrates the diversity in gene expression among different cancer types. Specifically, genes associated with BRCA are predominantly expressed at higher levels compared to those in BLCA, whereas ESCA-associated genes tend to have lower expression relative to the other cancer types examined. Additionally, a significant proportion of the top 100 genes identified by their LRP scores are implicated in transcription regulation and have associations with cancer, underscoring their potential roles in the molecular mechanisms underpinning BRCAness.

**Fig. 5.**
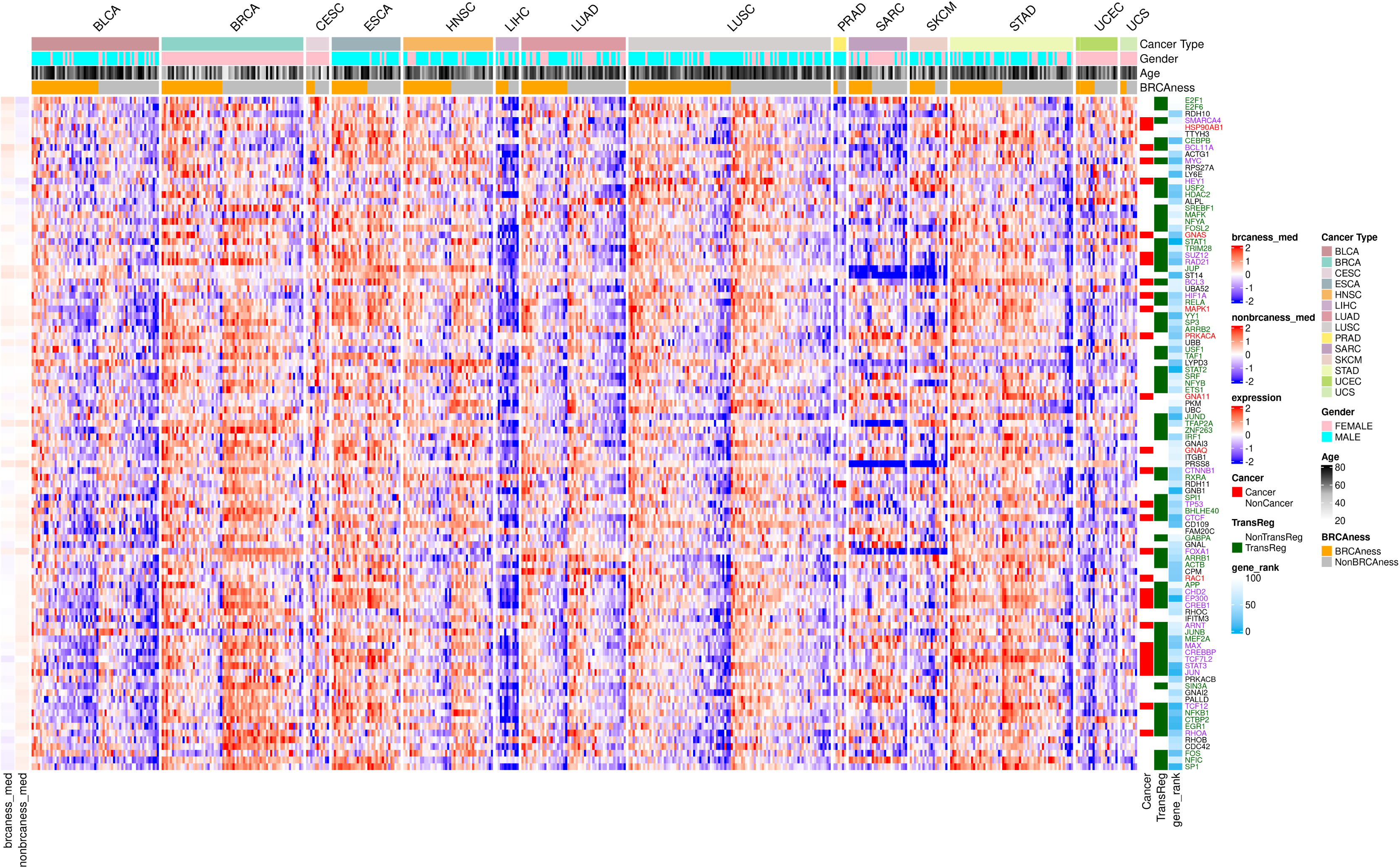
Comprehensive heatmap displaying the expression patterns of the top 100 genes with the highest LRP scores across multiple cancer types, distinguished by BRCAness and Non-BRCAness phenotypes. Each column represents a unique sample, while each row corresponds to a gene. The heatmap color spectrum, with red signifying elevated expression and blue indicating reduced expression, illustrates the differential gene activity within the two phenotypic classifications. Annotations above the heatmap categorize the samples by cancer type, gender, age, and BRCAness status, while side annotations identify genes recognized in the Cancer Gene Census (red), those involved in transcriptional regulation (green), and their relative LRP score-based ranks (blue).

The gene interaction network plot (Fig. 6) visualizes the functional interactions between genes that have top LRP scores, potentially highlighting their significance in the context of BRCAness. Each node in the network represents a gene, and the edges between nodes signify functional interactions between these genes. The color filling within the nodes denotes the differential median expression values between BRCAness and Non-BRCAness groups: a gradient from a deep color to a lighter shade could represent high to low expression differences, respectively. Shapes distinguish transcription regulators (boxes) from non-regulators (circles), suggesting potential regulatory roles within the network. Node borders colored red identify cancer-associated genes, while black signifies non-cancer genes. The colored regions group genes into relevant pathways or BRCAness-associated gene clusters, offering insights into their collective roles in the BRCAness phenotype. Among the top LRP genes within these pathway clusters, it is evident that two notable genes, TP53 and MYC, have been identified.

**Fig. 6.**
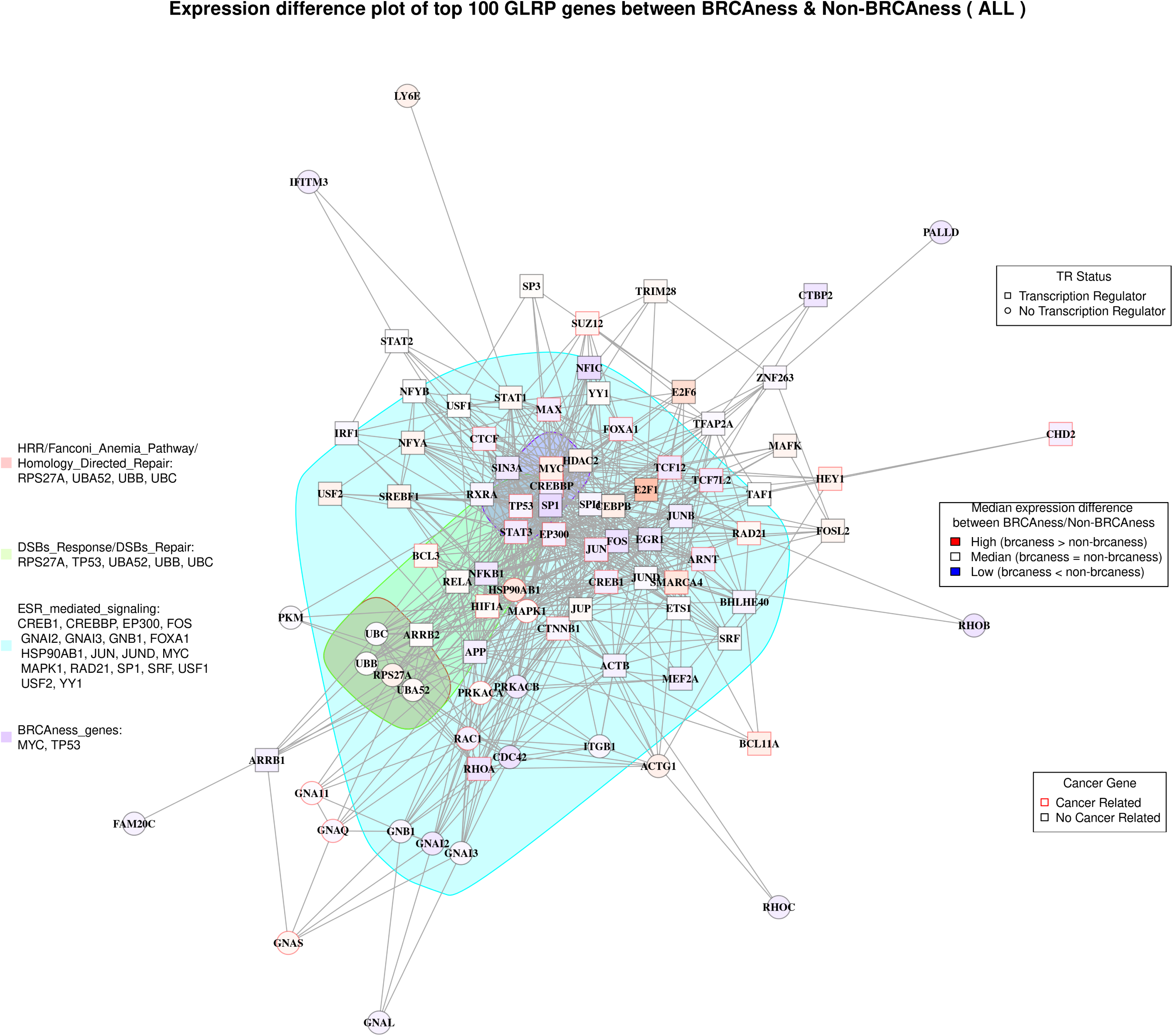
Gene interaction network plot illustrating the functional interplay among the top genes with the highest LRP scores in the context of BRCAness. Nodes represent individual genes, with the varying color fill indicating the difference in median expression between BRCAness and Non-BRCAness phenotypes. Box-shaped nodes highlight genes involved in transcription regulation, while circular nodes denote those without this function. Node border colors distinguish between cancer-associated genes (red) and non-cancer-related genes (black). The network is sectioned into colored regions, each grouping genes by their involvement in specific pathways or their association with BRCAness.

#### Comparative Analysis of Top Gene Contribution Patterns Between GLRP and RF Models in BRCA Data

Since feature importance analysis has been conducted in traditional machine learning models, the top 100 important genes were identified for each model. It was observed that only the top contributing genes from the Random Forest (RF) model intersect with genes associated with the BRCAness phenotype. Specifically, the genes AURKA and PLK1 were identified among the top 100 contributing genes in the RF model. Conversely, the important genes identified by other models, like Logistic Regression, Support Vector Classifier, and Decision Tree, did not include any genes implicated in the BRCAness. Thus RF will be selected to do the comparative analysis with GLRP.

Figurepresents a comparative analysis of gene expression heatmaps and interaction networks in BRCA cancer type, focusing on the important genes related to BRCAness identified by GLRP and RF. In the upper portion of the figure, the heatmaps delineate the expression patterns of the top 100 genes as determined by GLRP. These heatmaps reveal a pronounced involvement of genes associated with cancer and transcription regulation in the GLRP-selected set compared to the RF-selected set, with counts of 30 versus 4 for cancer-related genes, and 65 versus 26 for transcription regulation genes, respectively.

The network diagrams in the lower half of the figure further elucidate the biological pathways implicated by these genes. The GLRP-derived gene set prominently features genes involved in DNA repair and damage response pathways, as well as the Estrogen Receptor (ER) mediated signaling pathway, more so than the RF-derived set. The interaction network for the top GLRP genes illustrates a tightly-knit cluster of interactions, suggesting a strong functional relationship or co-regulation among genes associated with DNA repair or the BRCAness. In contrast, the network for the top RF genes displays a dispersed pattern, with significant genes being scattered and clusters more isolated. The nodes representing genes in the RF network are not as densely packed, and the interconnections are sparser, highlighting a less cohesive functional grouping.

**Fig. 7.**
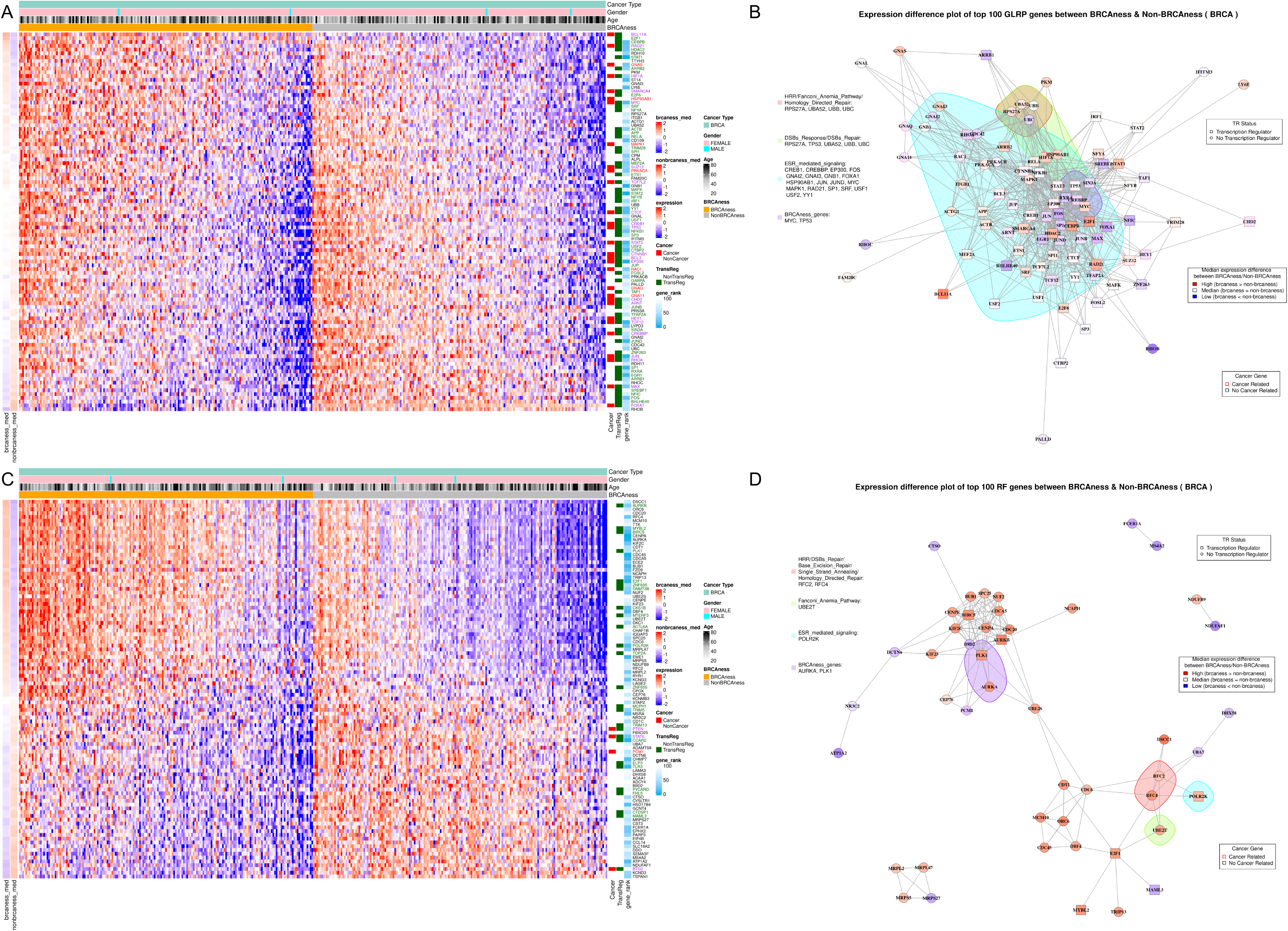
Gene importance comparaive analysis between RF and GLRP in BRCA dataset. A. The gene expression Heatmap of top 100 GLRP genes. B. The gene functional interaction network of top 100 GLRP genes. C. The gene expression Heatmap of top 100 RF genes. B. The gene functional interaction network of top 100 RF genes.

### 3.4 Comparative Analysis of Top GLRP Gene Contribution Patterns Between Cancer Types

Here, we conducted a comparative analysis of the top 100 GLRP gene expression and interaction network profiles across various cancer subtypes. Figure 8 presents a comparative analysis of the expression heatmaps and interaction networks for GLRP key genes associated with BRCAness in BRCA, OV, PAAD, and LUSC cancer types. Since the node color in the gene interaction network plot on the right represents the median difference in gene expression, it is readily apparent that this difference varies across different types of cancer. For instance, the TP53 gene exhibits almost no expression difference between the two groups in BRCA, but shows the most significant expression difference in OV, with higher expression in the Non-BRCAnes s group. The MYC gene is more highly expressed in the BRCAness group, whereas it is less expressed in OV. Additionally, in PAAD cancer type, genes involved in ESR-mediated signaling, such as EGR1 and FOS, are significantly less expressed in the BRCAness group than in the Non-BRCAness group. However, in lung squamous cell carcinoma (LUSC), the differences in expression of all top GLRP genes between the two groups are not pronounced.

**Fig. 8.**
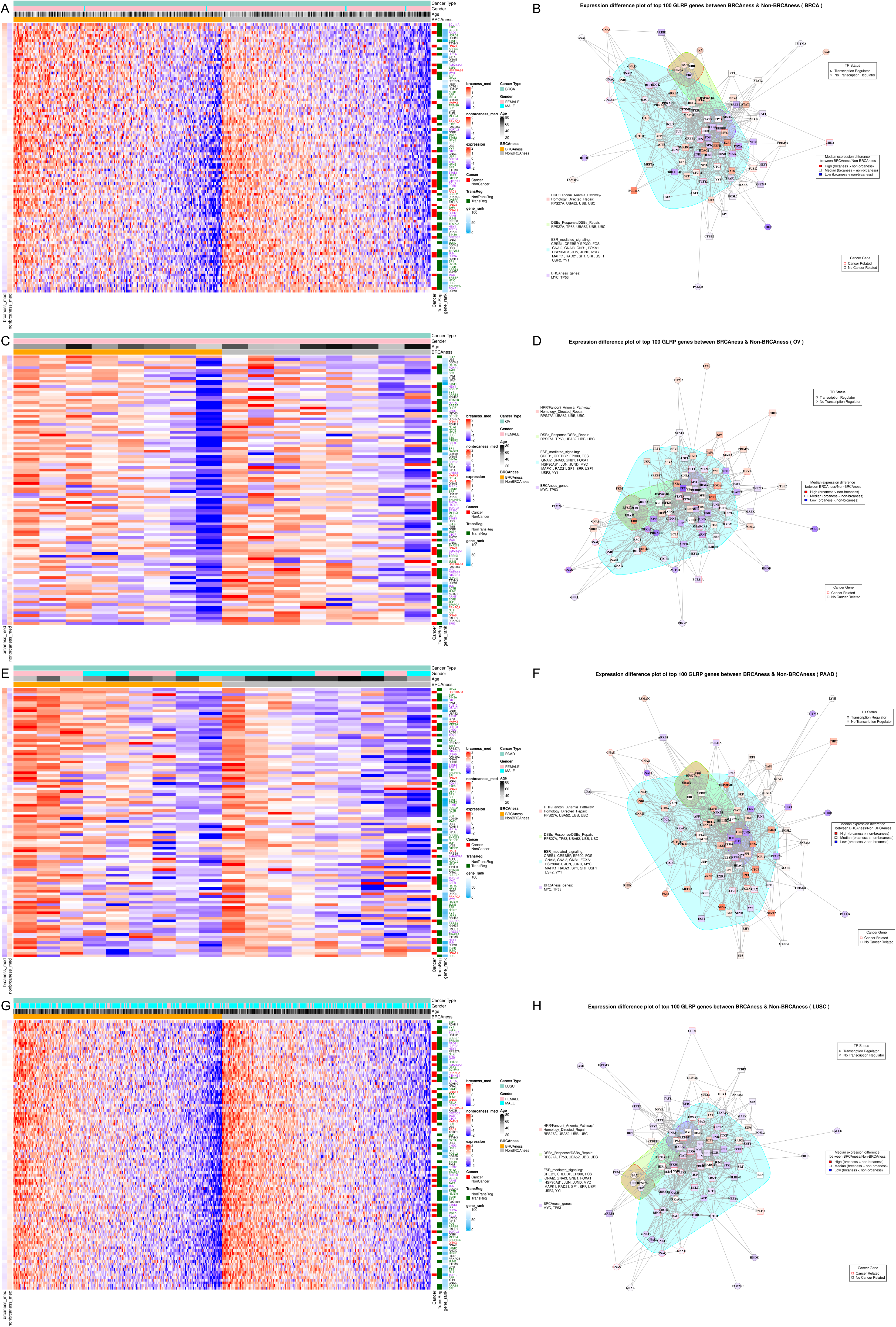
Gene importance comparative analysis between Cancer types. A. The gene expression Heatmap of top 100 GLRP genes in BRCA. B. The gene functional interaction network of top 100 GLRP genes in BRCA. C. The gene expression Heatmap of top 100 GLRP genes in OV. D. The gene functional interaction network of top 100 GLRP genes in OV. E. The gene expression Heatmap of top 100 GLRP genes in PAAD. F. The gene functional interaction network of top 100 GLRP genes in PAAD. G. The gene expression Heatmap of top 100 GLRP genes in LUSC. H. The gene functional interaction network of top 100 GLRP genes in LUSC.

Based on this observation, we selected the 20 GLRP genes with the most significant expression differences between the BRCAness and Non-BRCAness cohorts for each cancer type. We then conducted Gene Set Enrichment Analysis (GSEA) on these 20 differentially expressed genes across various cancer types to identify pathways that are highly associated with the BRCAness phenotype. Table 2 presents the consolidated results from the GSEA across multiple cancer types. Each row lists the ID and description of the pathways enriched based on these 20 differentially expressed genes, along with the number of cancer types in which each pathway was enriched and the specific cancer types where this enrichment was observed. The top-ranked pathways include ”Signaling by Nuclear Receptors,” ”Cellular Senescence,” ”Oncogene-Induced Senescence,” ”ESR-mediated Signaling,” and ”Parasitic Infection Pathways.”

**Table 2.** Summary table of the top 5 impacted pathways associated with the BRCAness phenotype across various cancer types in top 20 differential expressed LRP contributing genes GSEA analysis.

| ID | Description | Count | Cancer_type |
| --- | --- | --- | --- |
| R-HSA-9006931 | Signaling by Nuclear Receptors | 16 | ALL, BLCA, BRCA, CESC, COAD, ESCA, HNSC, LIHC, LUAD, OV, PAAD, READ, SARC, SKCM, UCEC, UCS |
| R-HSA-2559583 | Cellular Senescence | 15 | ALL, BLCA, BRCA, CESC, COAD, ESCA, HNSC, LUAD, LUSC, OV, PAAD, PRAD, READ, SARC, SKCM |
| R-HSA-2559585 | Oncogene Induced Senescence | 14 | ALL, BLCA, CESC, ESCA, HNSC, LUAD, LUSC, OV, PAAD, READ, SARC, SKCM, STAD, UCS |
| R-HSA-8939211 | ESR-mediated signaling | 13 | ALL, BLCA, CESC, COAD, ESCA, HNSC, LIHC, LUAD, PAAD, READ, SARC, SKCM, UCS |
| R-HSA-9824443 | Parasitic Infection Pathways | 13 | CESC, COAD, ESCA, HNSC, LIHC, LUSC, OV, PAAD, PRAD, READ, SARC, SKCM, UCS |

## 4 Discussion

Our study presents a comprehensive analysis of BRCAness phenotype across various cancer types, utilizing graph layer-wise propagation (GLRP) as well as traditional methods such as Random Forest (RF) and DGE analysis. We have uncovered significant insights into the genetic underpinnings of the BRCAness, which is crucial for understanding its role in cancer development and progression, and potentially guiding personalized treatment strategies.

### 4.1 Limitations of DGE Analysis in BRCAness Phenotype Interpretation

Differential gene expression (DGE) analysis has been a cornerstone in identifying genes associated with various phenotypes, including BRCAness. However, our findings highlight its limitations, primarily due to compensatory mechanisms, the disparity between gene expression and functionality, and intragroup heterogeneity. Unlike DGE, GLRP leverages a gene functional interaction network, offering a more nuanced interpretation of BRCAness. This method not only identifies genes directly involved in BRCAness and DNA repair but also elucidates the complex biological interactions underpinning this phenotype. The network-based approach of GLRP allows for the identification of genes that, while not differentially expressed, play critical roles in the BRCAness phenotype through their functional interactions and pathways.

### 4.2 Comparative Analysis of Feature Importance in RF and GLRP

In the comparative analysis of feature importance between Random Forest (RF) and Graph Layer-wise Relevance Propagation (GLRP), it becomes evident that each method offers unique insights into gene significance, albeit from different perspectives. RF gauges feature importance based on the reduction of impurity within decision trees, such as through Gini impurity or entropy measures. This approach prioritizes genes with the most significant impact on predictive accuracy, potentially overlooking those with subtle yet crucial biological roles in cancer. Consequently, RF may not always capture genes that are biologically significant but do not markedly enhance classification performance across its ensemble of trees. Conversely, GLRP transcends this limitation by integrating gene expression data with a gene functional interaction network, thus grounding feature importance in established biological relationships. This network-centric approach enables GLRP to recognize the importance of genes not merely through their expression levels but by their roles and interconnections within the network. As a result, GLRP is particularly effective in pinpointing genes pivotal to pathways like DNA repair and Estrogen Receptor (ER)-mediated signaling, both of which are intricately linked to the BRCAness. DNA repair pathway deficiencies reflect the hallmark feature of BRCAness, making the identification of genes within these pathways crucial for understanding and targeting this phenotype [37]. Similarly, alterations in ER-mediated signaling pathways have been implicated in the development and therapeutic responses of BRCAness-affected tumors [38], further highlighting the relevance of these pathways to the phenotype. GLRP’s ability to elucidate the inter-connected and functionally significant genes within these critical pathways provides a deeper, more biologically nuanced perspective on BRCAness. This approach not only enhances our understanding of the genetic underpinnings of BRCAness but also underscores GLRP’s superior capability in unraveling the complex genomic landscape associated with this unique cancer phenotype, as demonstrated by studies highlighting the modulation of BRCA1-mediated DNA damage repair by deregulated ER-*α* signaling in breast cancers [37] and the crucial role of estrogen signaling in the DNA damage response in hormone-dependent breast cancers [39]. Furthermore, the phenomenon of BRCA1-mutated estrogen receptor-positive breast cancer showing BRCAness suggests sensitivity to drugs targeting homologous recombination deficiency, emphasizing the importance of these pathways in the context of therapeutic intervention [40].

### 4.3 Insights from Top GLRP Genes: Transcription Regulation and Cancer-Related Associations

In the analysis of the top 100 genes identified by GLRP, a significant emphasis is placed on transcription regulation and direct cancer-related associations, underscoring the intricate role of transcriptional control in BRCAness. A remarkable 65% of these genes are implicated in transcription regulation processes, highlighting the critical influence of genes such as BCL11A, RAD21, STAT3, and E2F1. These genes, known for their fundamental roles in cell cycle control, apoptosis, and DNA repair, suggest the presence of a vital regulatory network potentially disrupted in BRCAness [41–47]. This disruption could be a driving factor behind the phenotype’s hallmark traits, including genomic instability and enhanced sensitivity to DNA-damaging therapies. Furthermore, the identification of 30 genes directly tied to cancer processes within the top GLRP genes underscores the immediate role these genes play in tumorigenesis and BRCAness-specific pathways. Among these, the inclusion of pivotal genes like TP53 and MYC, integral to cell growth and genomic integrity, indicates their direct contribution to the BRCAness phenotype [48–50]. This connection offers insights into tumor behavior, response to treatment, and prognosis, painting a comprehensive picture of the genetic landscape pivotal to BRCAness and its underlying mechanisms.

### 4.4 Elucidating BRCAness Through GSEA Pathway Analysis Across Cancer Types

The comparison of gene interaction network plots across different cancers reveals notable variance in the median expression differences of genes, underscoring the heterogeneous nature of BRCAness. This variance emphasizes the adaptive and context-dependent expression of genes associated with BRCAness, suggesting that the phenotype’s manifestation is intricately connected to the specific biological landscape of each cancer type. By extracting the top 20 genes with the most significant median expression differences among the 100 most impactful GLRP genes for each cancer type and applying Gene Set Enrichment Analysis (GSEA), this study illuminates the critical pathways that recur across cancers, directly linking them to BRCAness and DNA repair mechanisms. This analysis highlights the critical roles of ”Signaling by Nuclear Receptors,” ”Cellular Senescence,” ”Oncogene-Induced Senescence,” ”Estrogen Receptor (ESR)-mediated signaling,” and ”Parasitic Infection Pathways” in the underlying mechanisms that define BRCAness, drawing a comprehensive picture of its biological foundations. These pathways, integral to cellular regulation, stress response, and genomic integrity, offer profound insights into the molecular intricacies that underpin the BRCAness phenotype. They suggest a multifaceted relationship with DNA repair and cellular resilience mechanisms, further elucidating the complexity of BRCAness across diverse cancer types.

Signaling by Nuclear Receptors serves as a cornerstone for hormone-mediated cellular responses, crucial for the maintenance of cellular homeostasis and the regulation of gene expression [51]. In the context of BRCAness, aberrations in this pathway can lead to a dysregulated response to hormonal signals, potentially influencing the efficacy of DNA repair mechanisms and impacting cellular sensitivity to DNA-damaging agents [51, 52]. This pathway’s involvement underscores the complex interplay between hormonal signaling and genomic stability, highlighting its potential as a therapeutic target in cancers exhibiting BRCAness characteristics.

Cellular Senescence and Oncogene Induced Senescence pathways represent critical cellular safeguards against uncontrolled proliferation and tumorigenesis. These pathways, activated in response to DNA damage or oncogenic stress, halt cell cycle progression, providing a window for DNA repair or triggering apoptosis in irreparably damaged cells [53–55]. The enrichment of these pathways in BRCAness-associated cancers suggests a heightened cellular effort to maintain genomic integrity in the face of defective DNA repair capabilities [55, 56]. This insight into the senescence mechanisms offers a unique perspective on the adaptive strategies employed by cells to counterbalance the vulnerabilities introduced by BRCAness, including the potential for therapeutic intervention that exploits these pathways.

ESR-mediated signaling is pivotal in mediating the effects of estrogens on cellular growth, differentiation, and survival [57, 58]. The association of this pathway with BRCAness points to the critical role of estrogen signaling in modulating the DNA damage response and repair processes [39, 59]. Disruptions in ESR-mediated signaling in BRCAness may contribute to the phenotype’s sensitivity to PARP inhibitors and other therapies targeting DNA repair pathways, suggesting a hormone-dependent dimension to the genomic instability characteristic of BRCAness [60].

Lastly, the Parasitic Infection Pathway highlights an unexpected connection between infectious agents and cancer, reflecting the broader context of cellular stress responses. While seemingly unrelated, the activation of these pathways in BRCAness-associated cancers could indicate a general upregulation of cellular defense mechanisms against various forms of stress, including genomic instability. This association invites further investigation into the role of immune and stress response pathways in the manifestation of BRCAness, potentially uncovering novel intersections between infection, immunity, and cancer biology that are relevant to BRCAness [61, 62].

## 5 Conclusion

In this comprehensive study, we elucidated the BRCAness phenotype across various cancer types, showcasing the advantages of employing Graph Layer-wise Relevance Propagation (GLRP) over traditional Differential Gene Expression (DGE) analysis and machine learning methods like Random Forest (RF). GLRP provided a deeper understanding of the BRCAness phenotype, enabling the identification of crucial genes within key biological pathways, such as DNA repair and Estrogen Receptor (ESR)-mediated signaling, which are not as readily apparent through DGE or conventional ML approaches. A significant portion of the top genes identified by GLRP is involved in transcription regulation (65%) and directly linked to cancer processes (30%), underlining the complexity and importance of transcriptional control and specific oncogenes in the manifestation of BRCAness. Moreover, through Gene Set Enrichment Analysis (GSEA) of the top differentially expressed genes, our study identifies pivotal pathways, including Signaling by Nuclear Receptors, Cellular Senescence, and ESR-mediated signaling, across cancer types, elucidating their potential roles in BRCAness and DNA repair mechanisms. These insights not only enhance our understanding of the genetic and molecular foundations of BRCAness but also highlight the potential for developing more targeted and personalized therapeutic strategies for cancer patients exhibiting this complex phenotype.

## Supplementary information

- **Supplementary File S1. Gene interaction network plots and heatmaps by cancer types.**This file includes three types of files:

– XXX Network Plot.png: Gene interaction network plot for a specific cancer type.
– XXX Heatmap.png: Heatmap displaying the expression patterns of the top 100 genes with the highest LRP scores for a specific cancer type.
– XXX Pathway result.csv: Gene set enrichment analysis results for the top 20 DE genes among the top 100 GLRP score genes for a specific cancer type.
- **Supplementary Table S1. Top 100 genes by LRP scores.**
- **Supplementary Table S2. Summary results of gene set enrichment analysis across multiple cancer types.**

## Acknowledgements

We would like to acknowledge the Volkswagen Foundation’s support in MTB-report project. JY was supported by the Ph.D. program ”Genome Science” - International Max Plank Research School, part of the Göttingen Graduate Center for Neurosciences, Biophysics, and Molecular Biosciences. We acknowledge the use of ChatGPT, for assisting in refining the manuscript text. The model was employed to enhance clarity, improve grammar, and suggest revisions based on the author’s input. All final decisions regarding the content and structure of the manuscript were made by the authors.

## Declarations

### Conflict of interest/Competing interests (check journal-specific guidelines for which heading to use)

The authors declare no conflict of interest.

### Ethics approval and consent to participate

Not applicable

### Consent for publication

Not applicable.

### Code availability

https://gitlab.gwdg.de/MedBioinf/mtb/brcaness_glrp_deciphering.

### Author contribution

JY, JD, and TB conceived and designed the study. JY collected data and designed specific analysis details. JD and TB provided insights into molecular biology and suggestions for analysis improvements. HC provides insights and assistance on machine learning, such as GCNN and LRP. JY wrote the manuscript, while JD, AB and TB were responsible for editing the manuscript.

