## Supplementary figures and images for "Deciphering BRCAness Phenotype in Cancer: A Graph Convolutional Neural Network Approach with Layer-wise Relevance Propagation Analysis"

### ALL_CancerType_Split_Heatmap.png

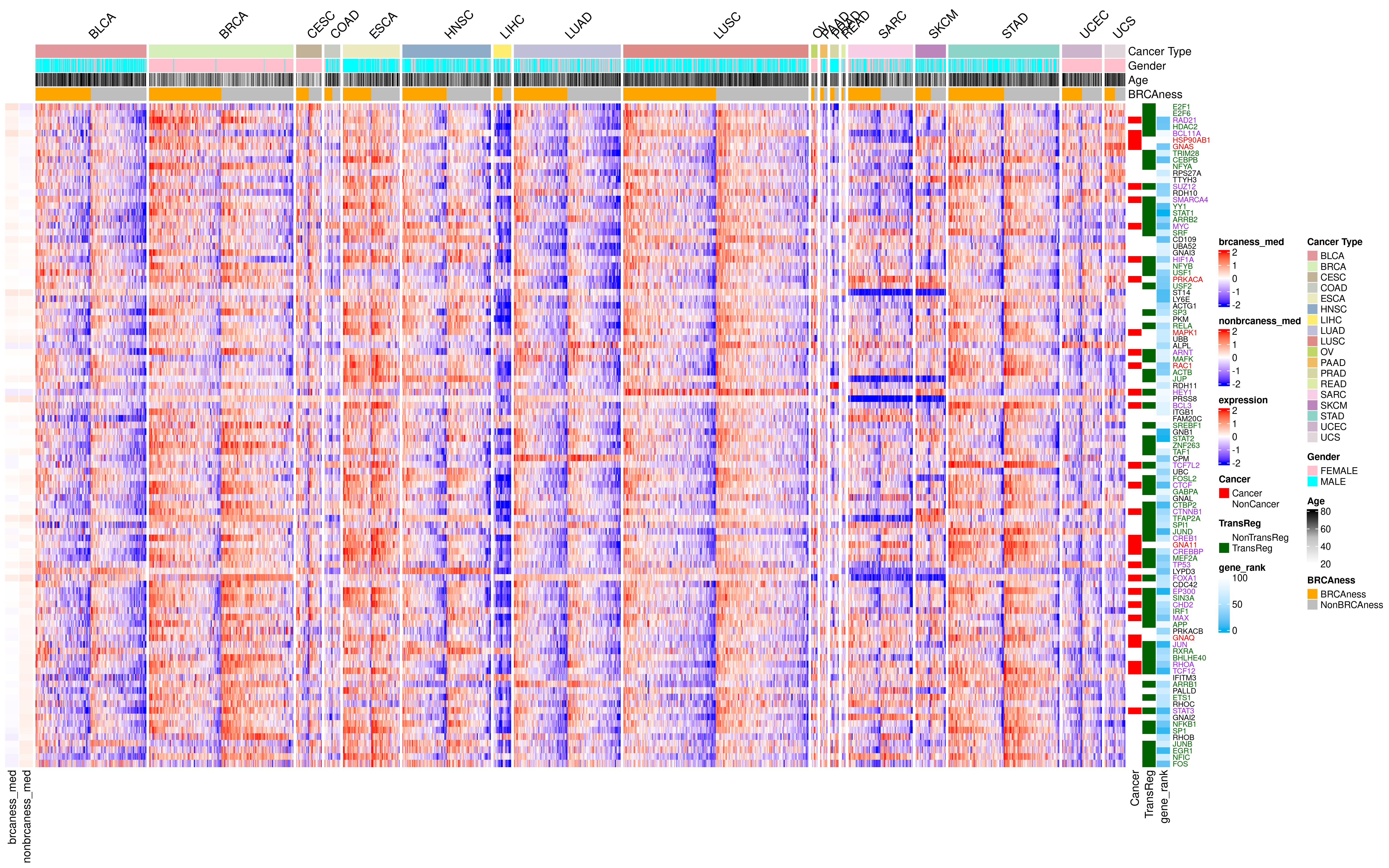

### ALL_Heatmap.png

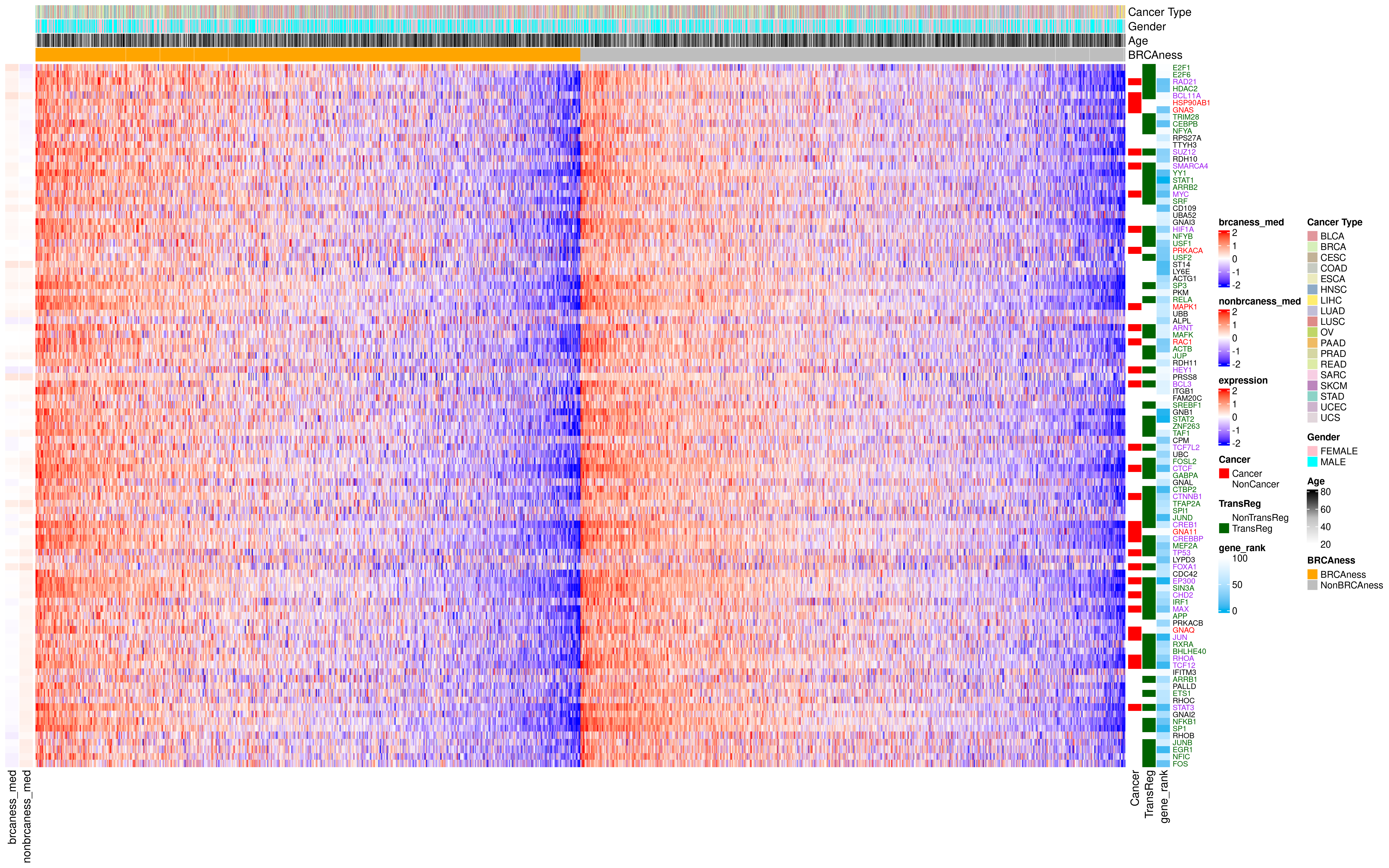

### ALL_Network_Plot.png

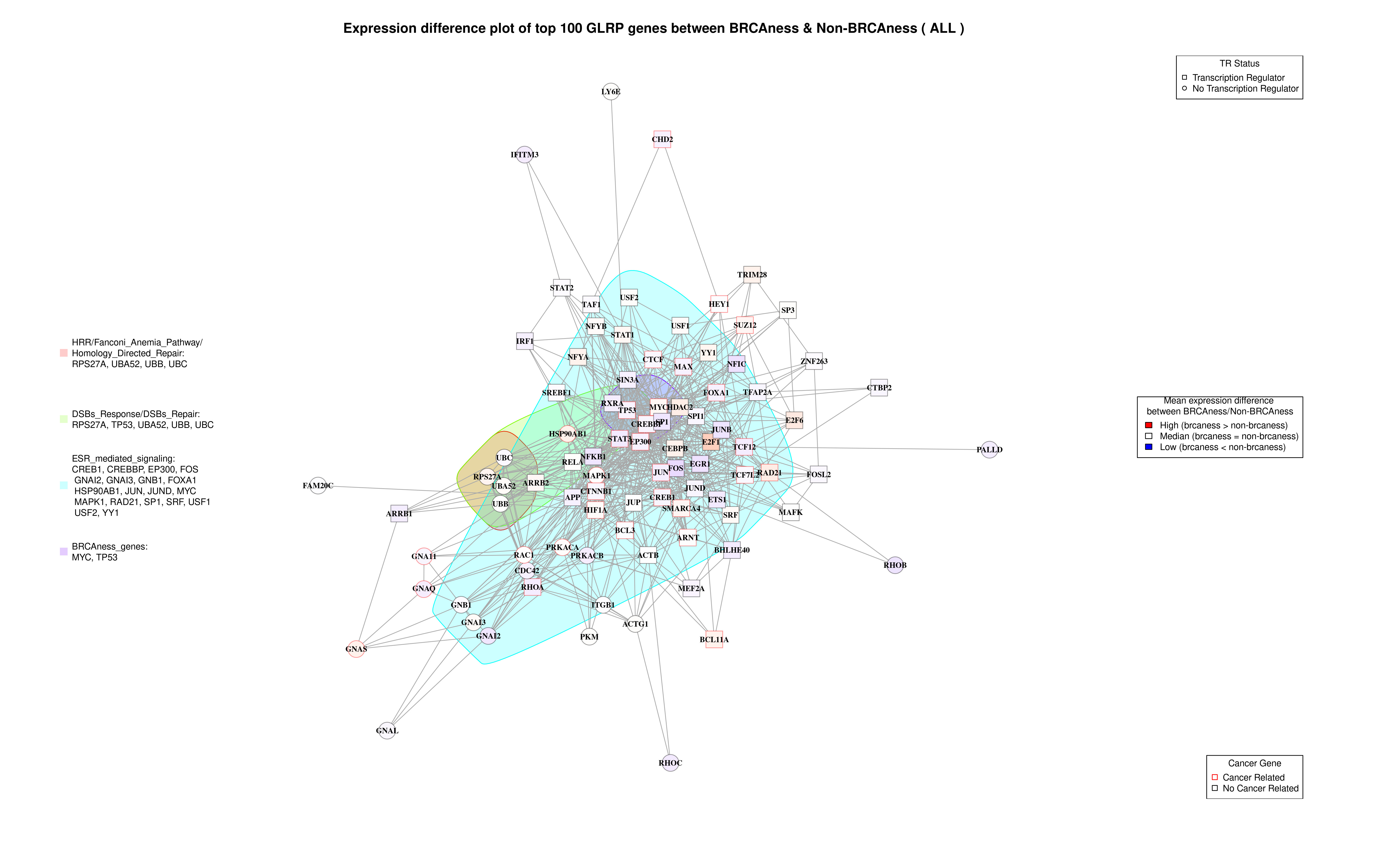

### BLCA_Heatmap.png

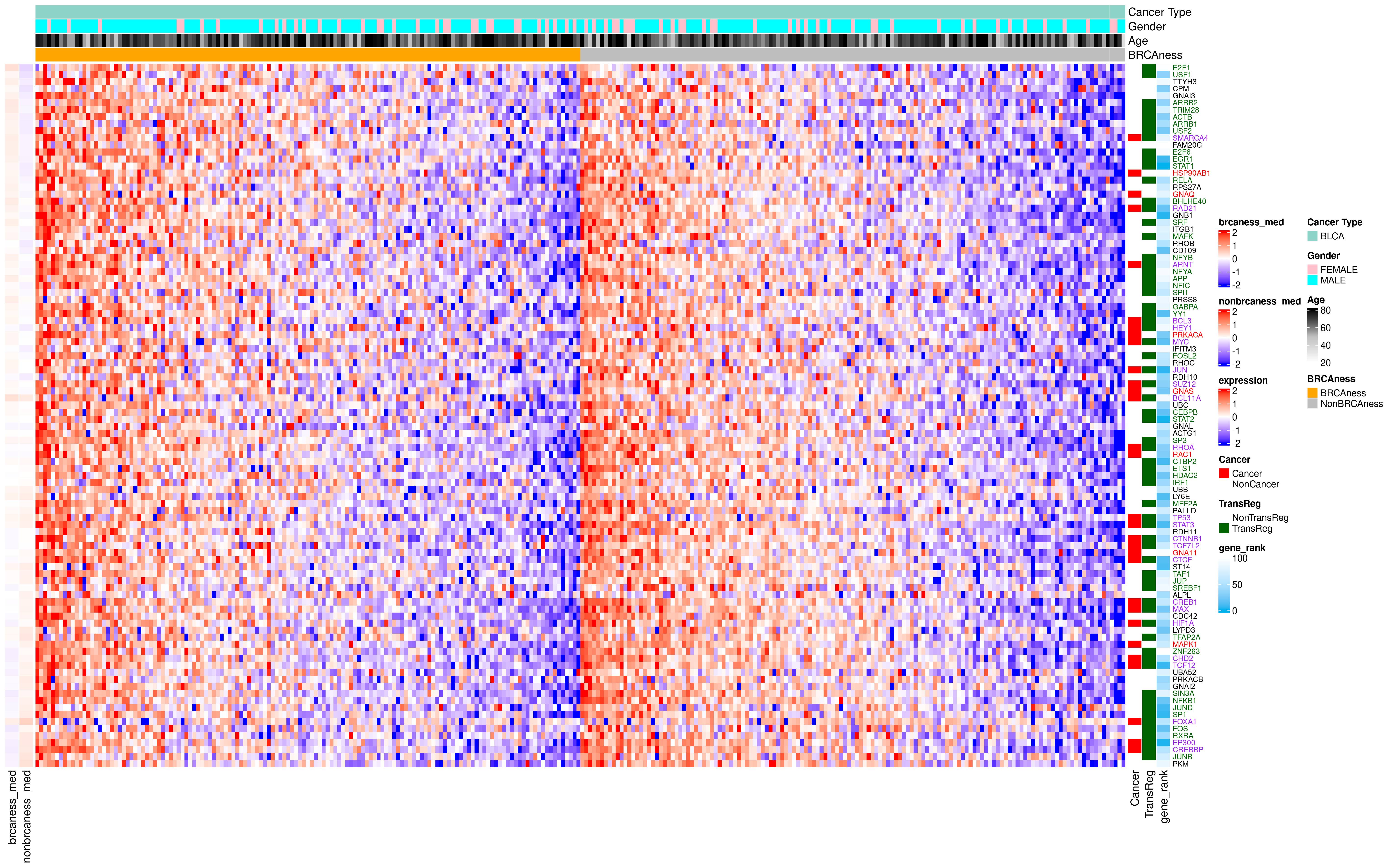

### BLCA_Network_Plot.png

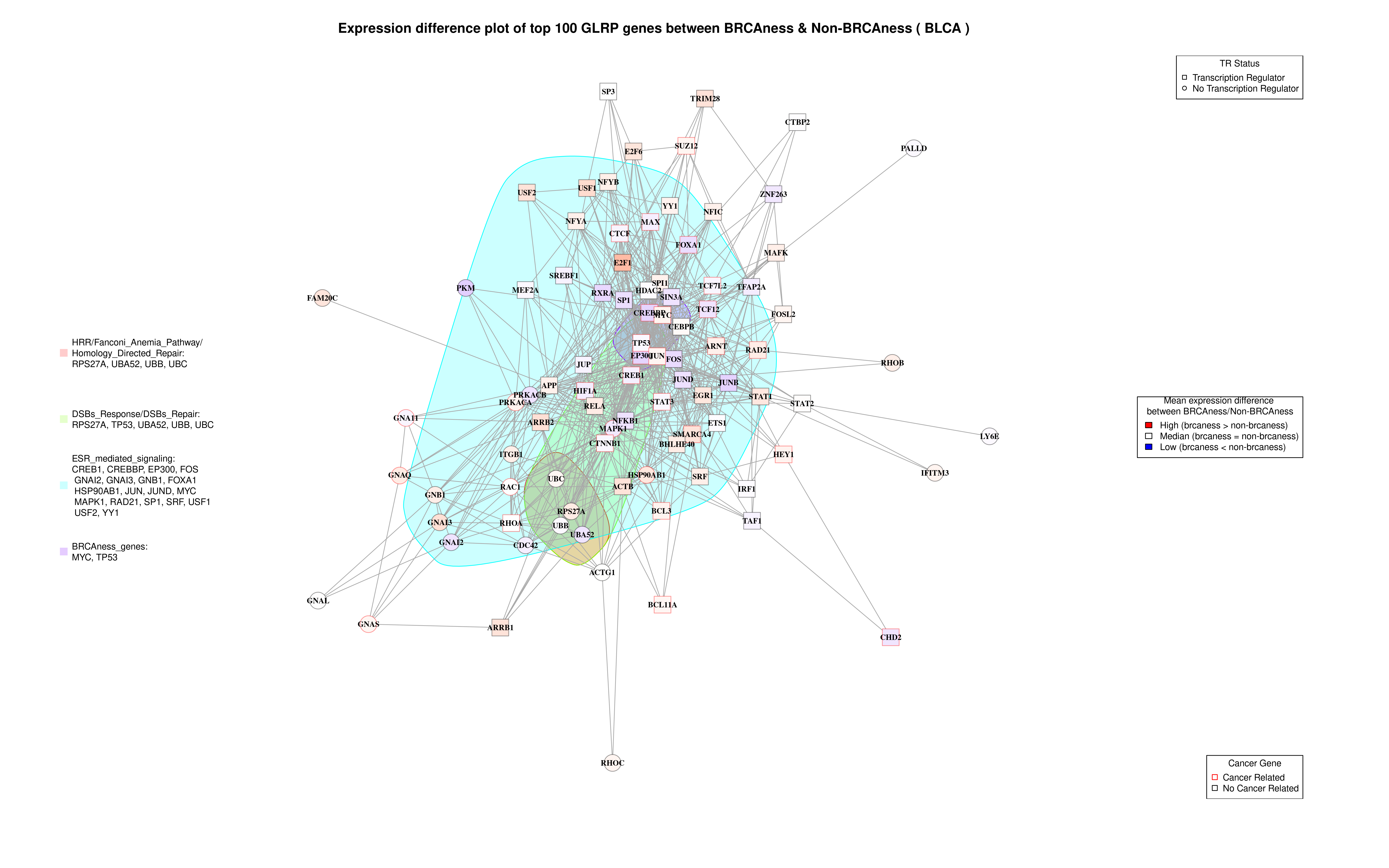

### BRCA_Heatmap.png

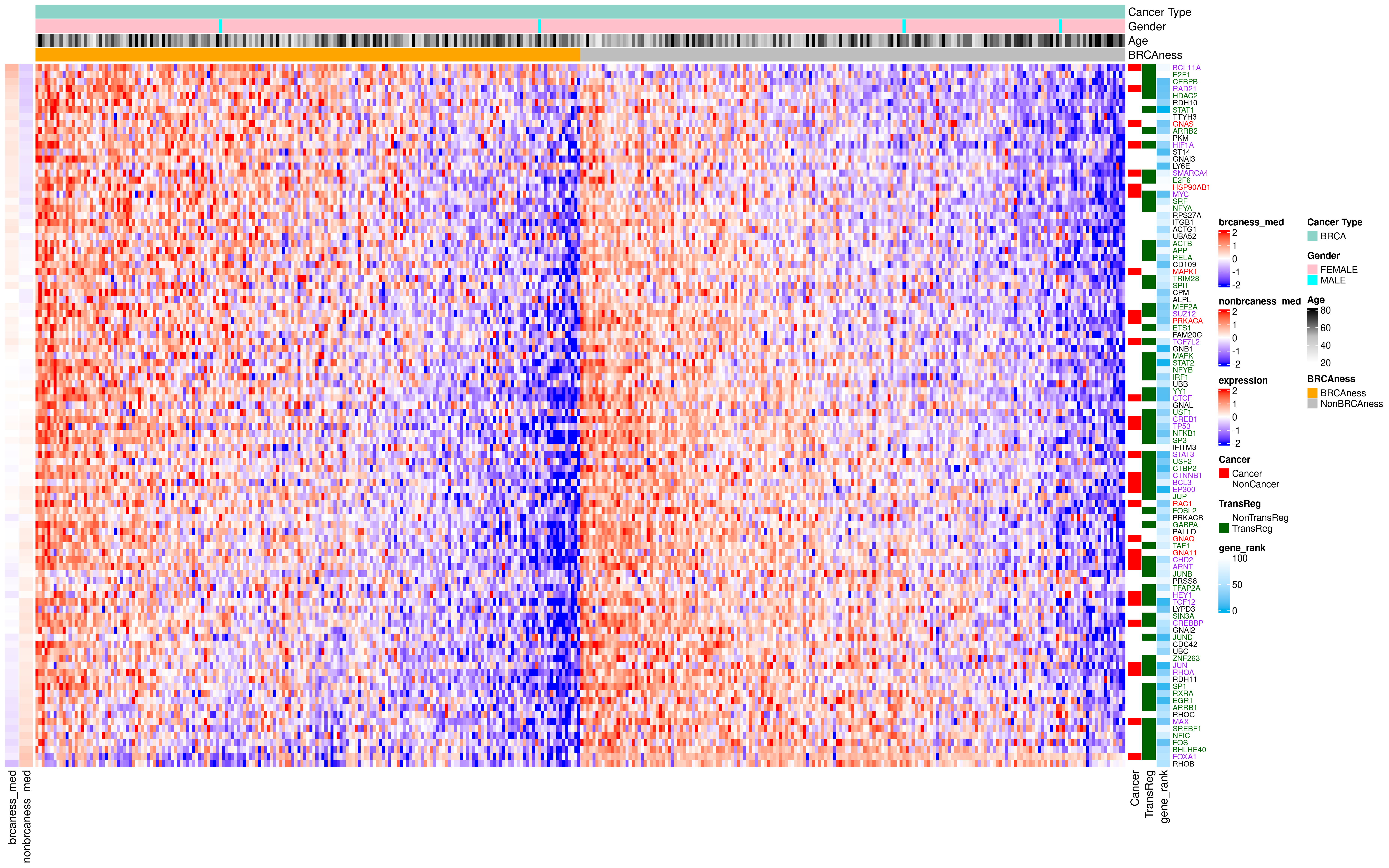

### BRCA_Network_Plot.png

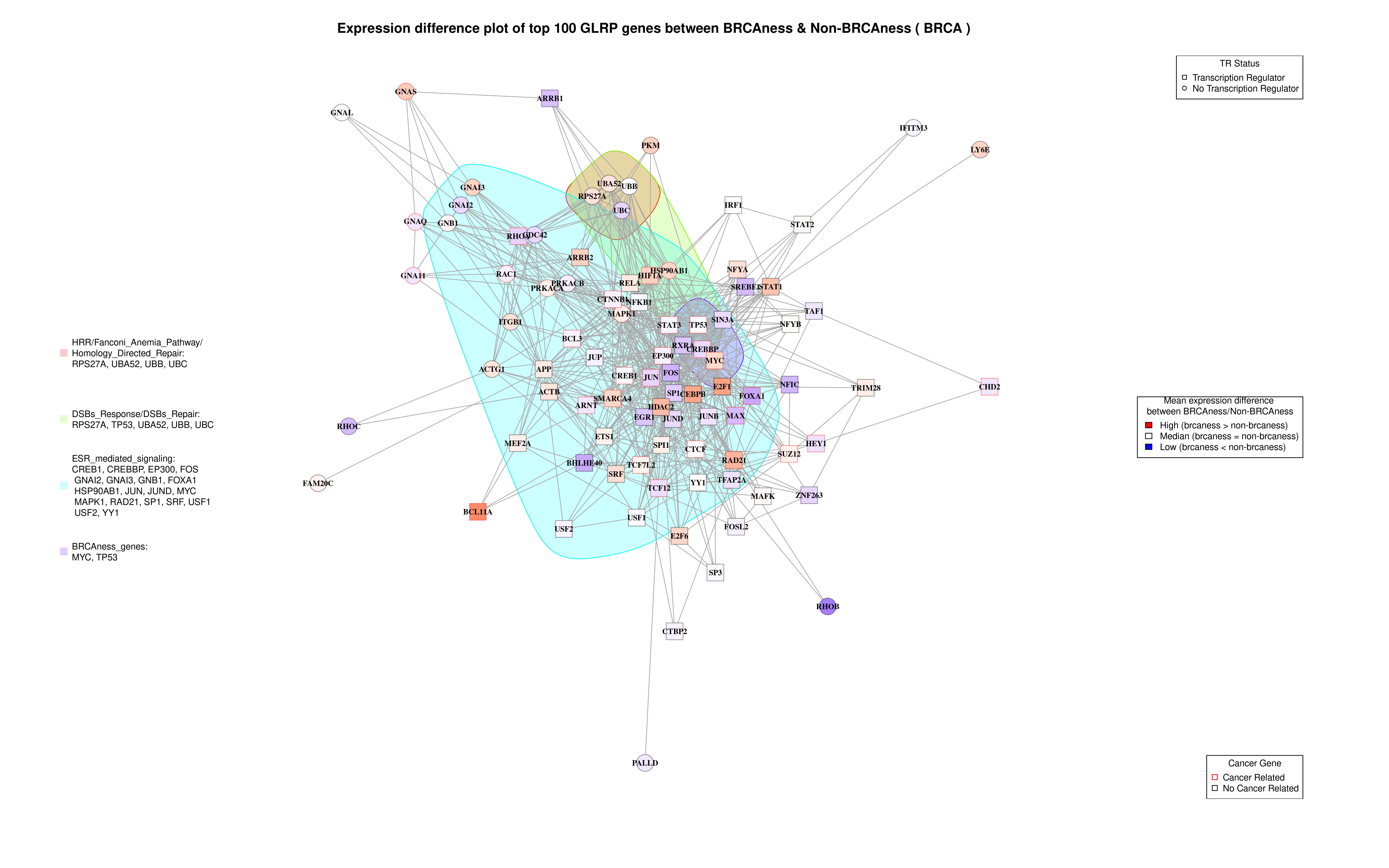

### CESC_Heatmap.png

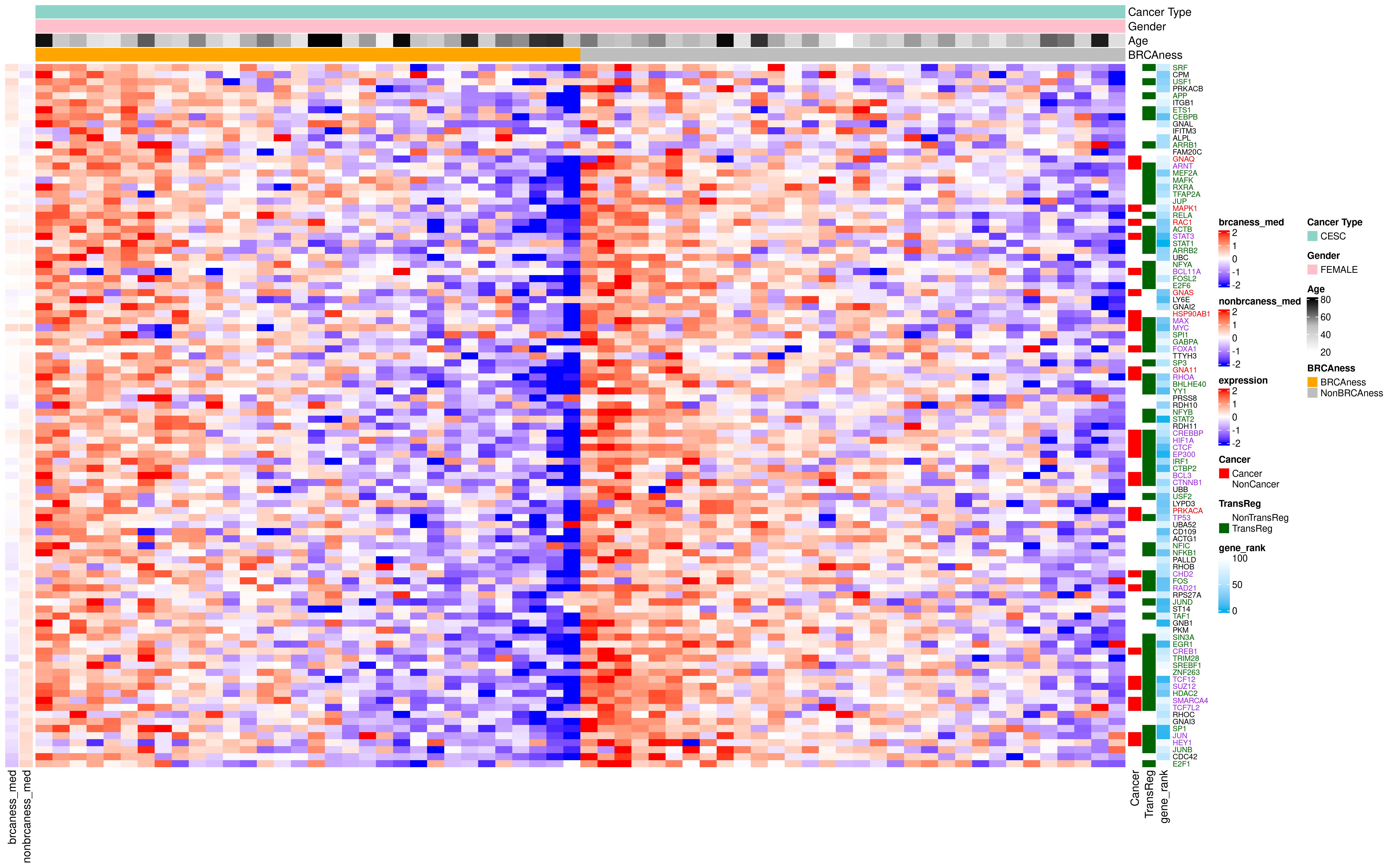

### CESC_Network_Plot.png

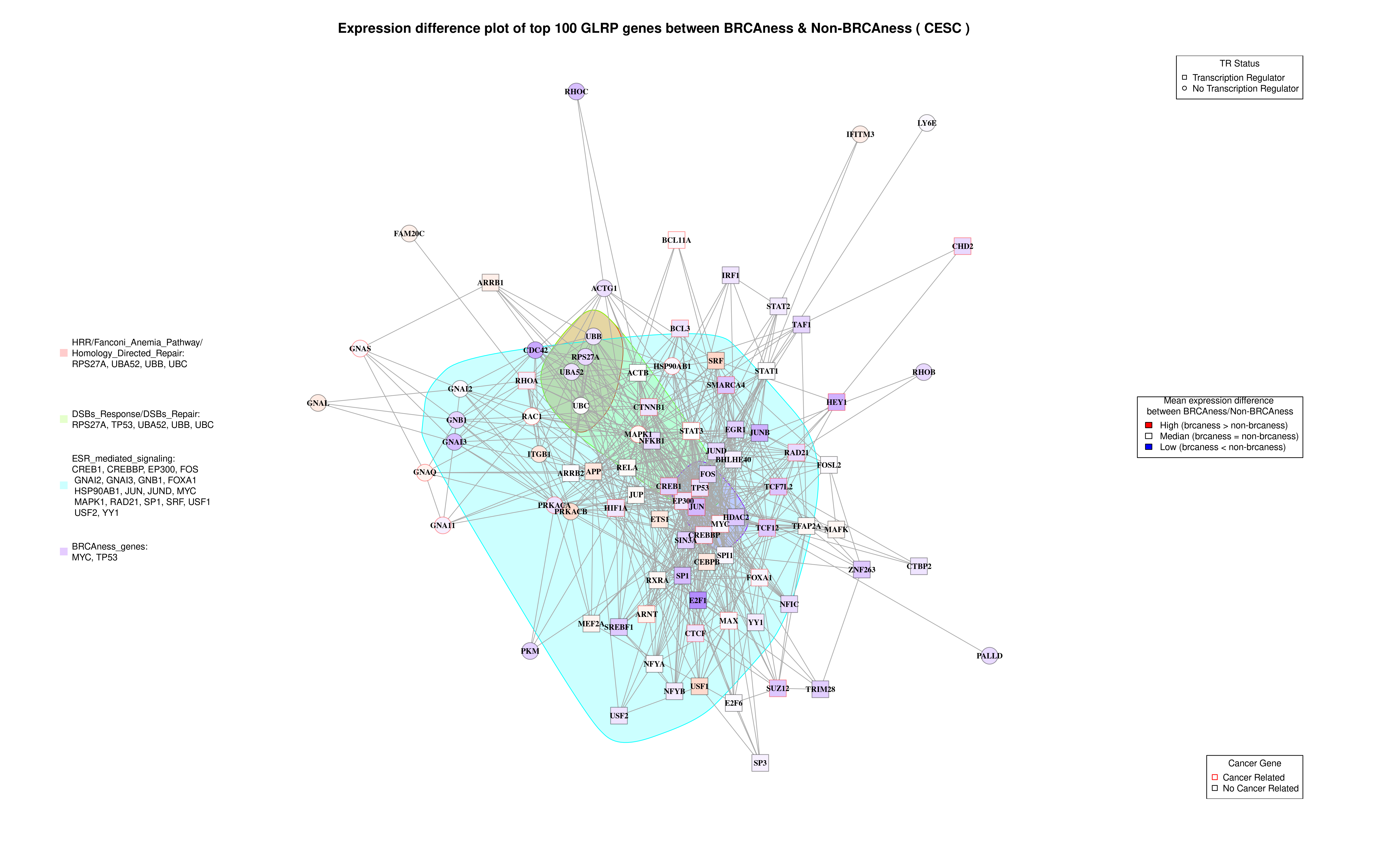

### COAD_Heatmap.png

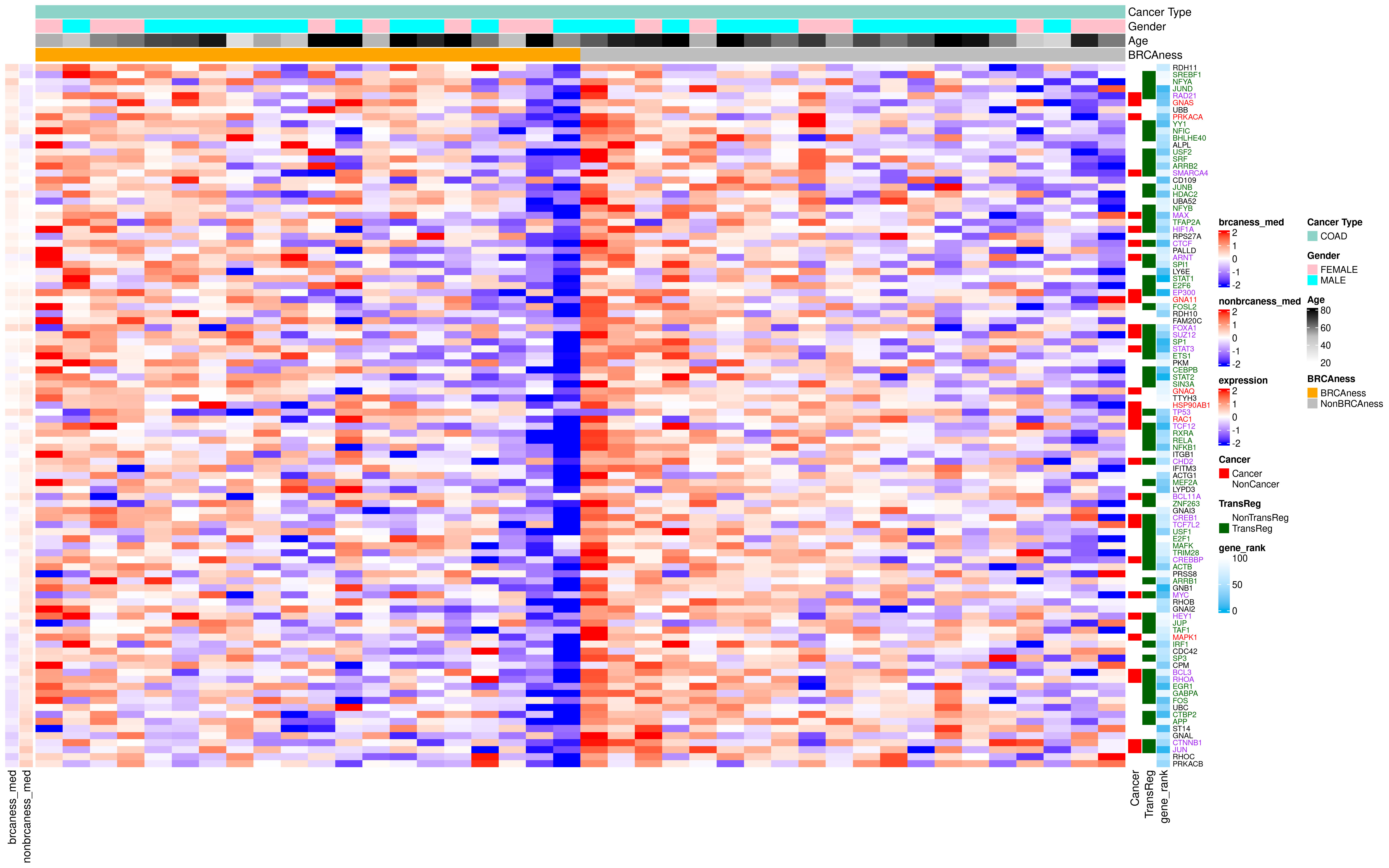

### COAD_Network_Plot.png

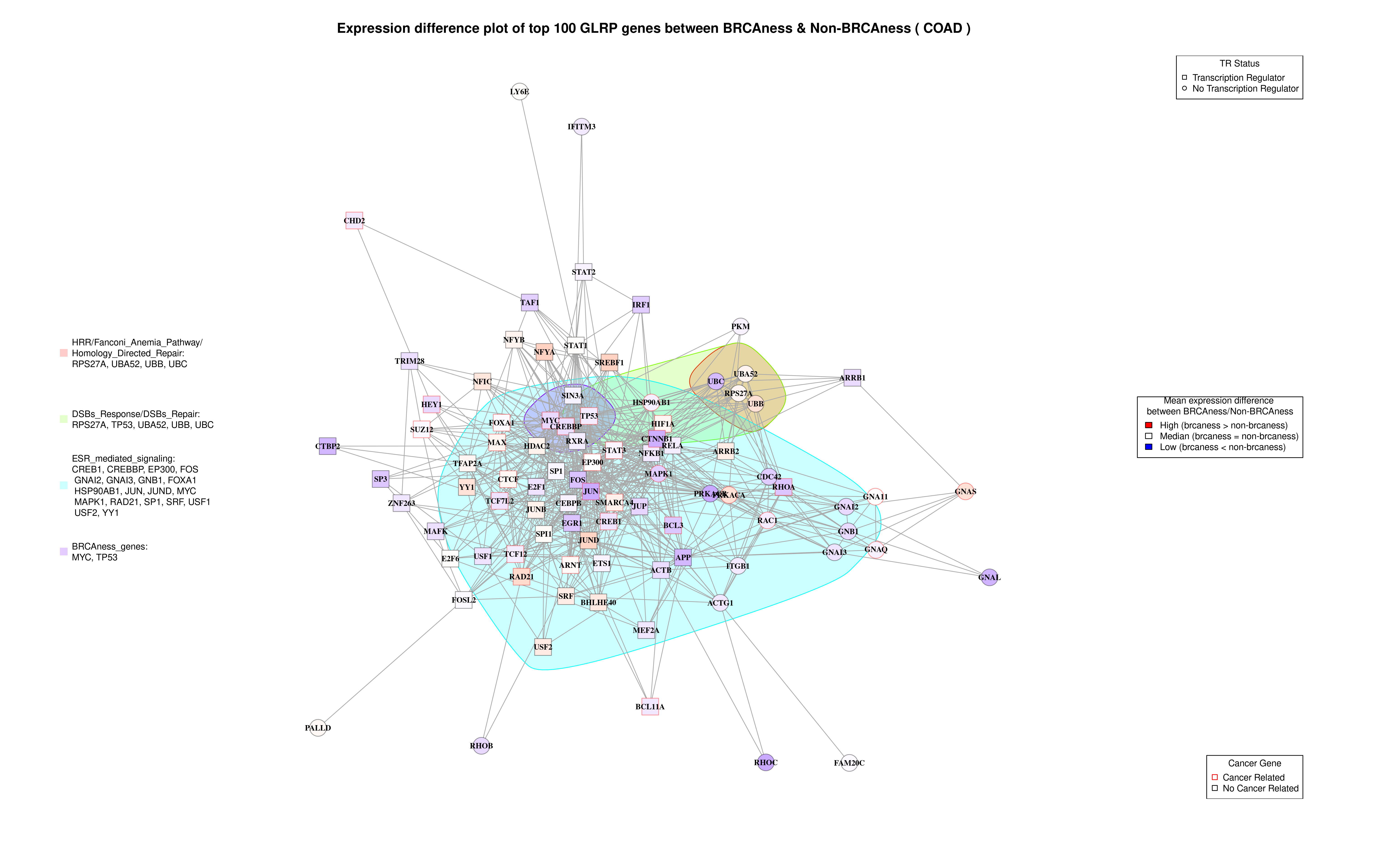

### ESCA_Heatmap.png

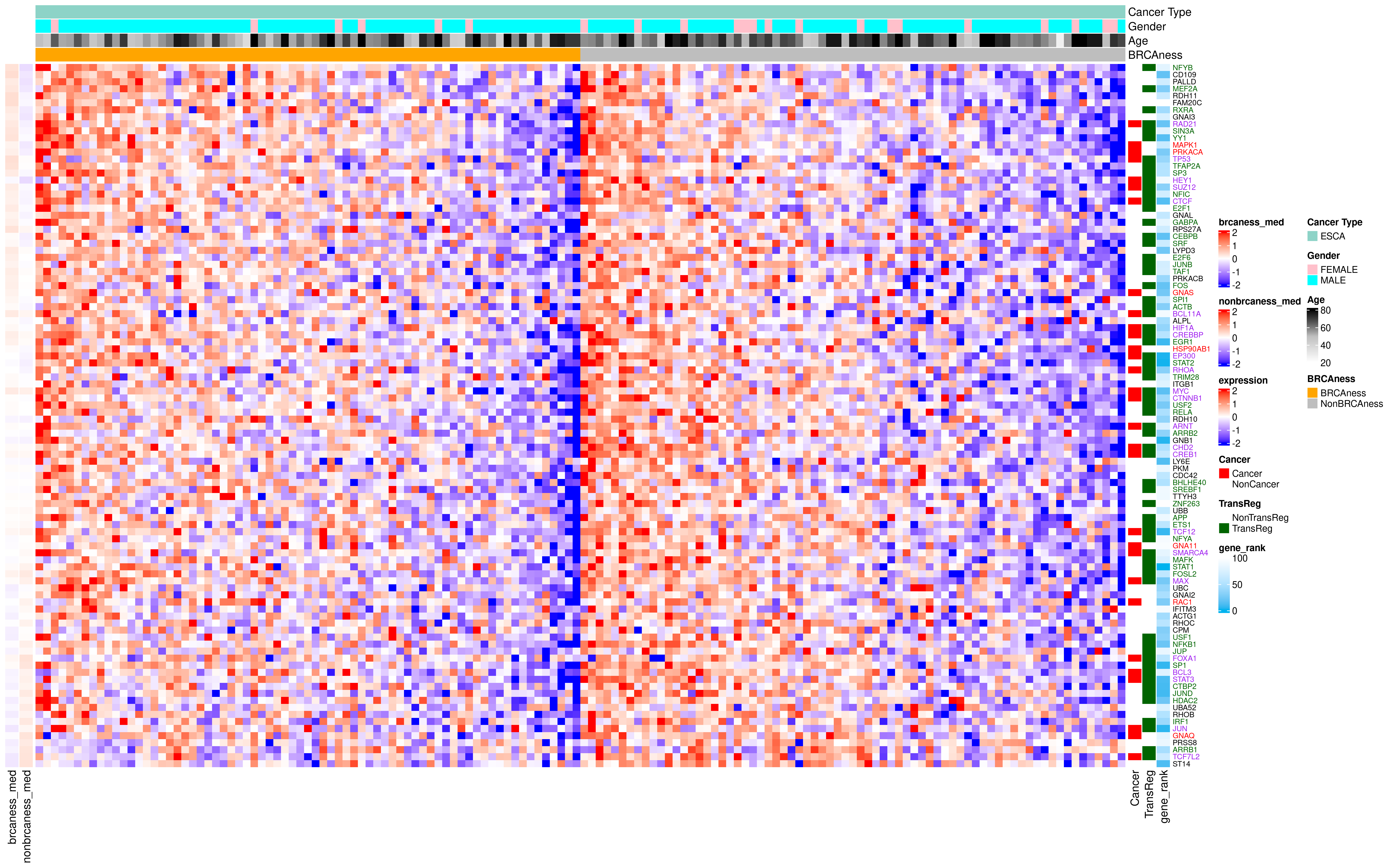

### ESCA_Network_Plot.png

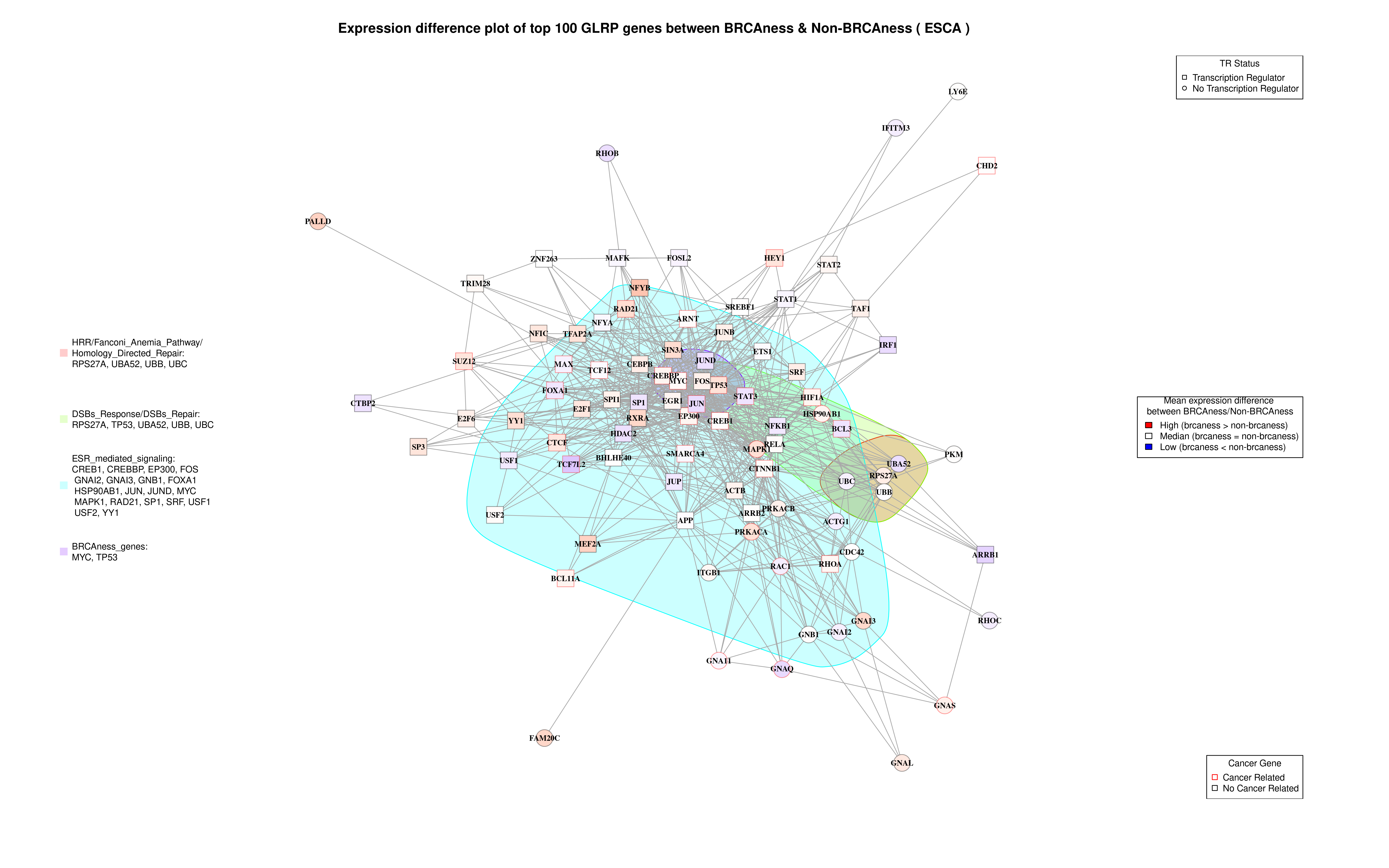

### HNSC_Heatmap.png

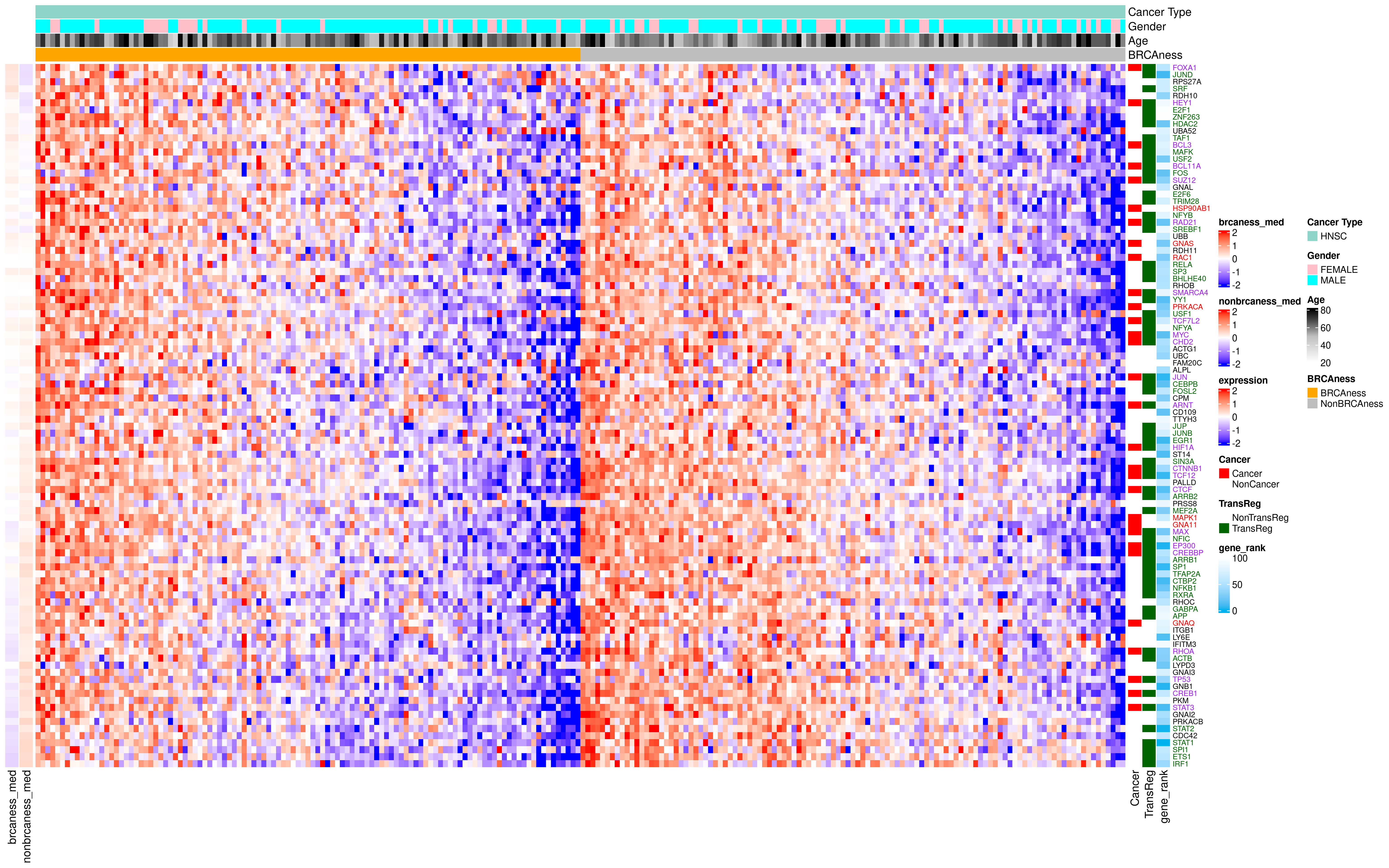

### HNSC_Network_Plot.png

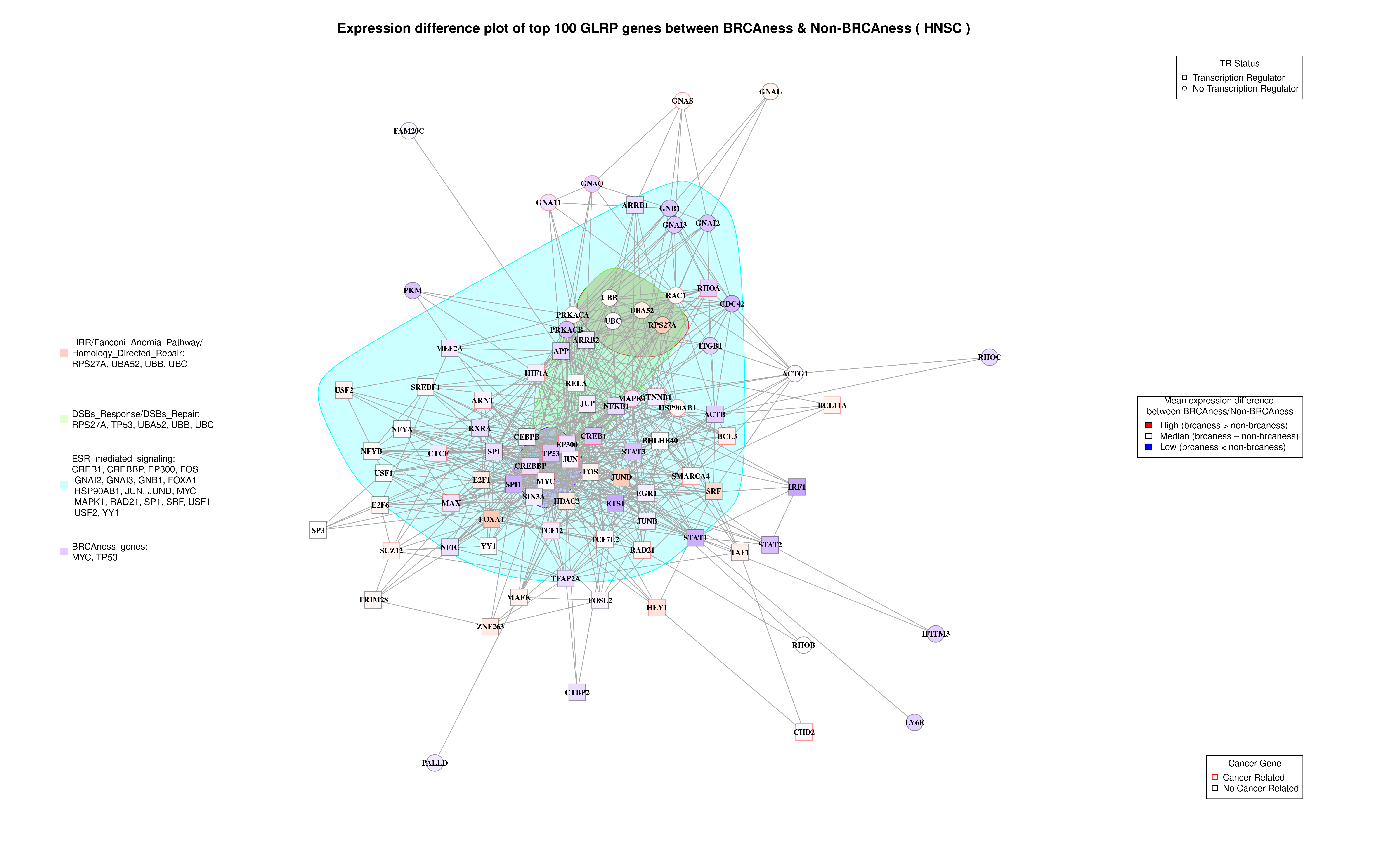

### LIHC_Heatmap.png

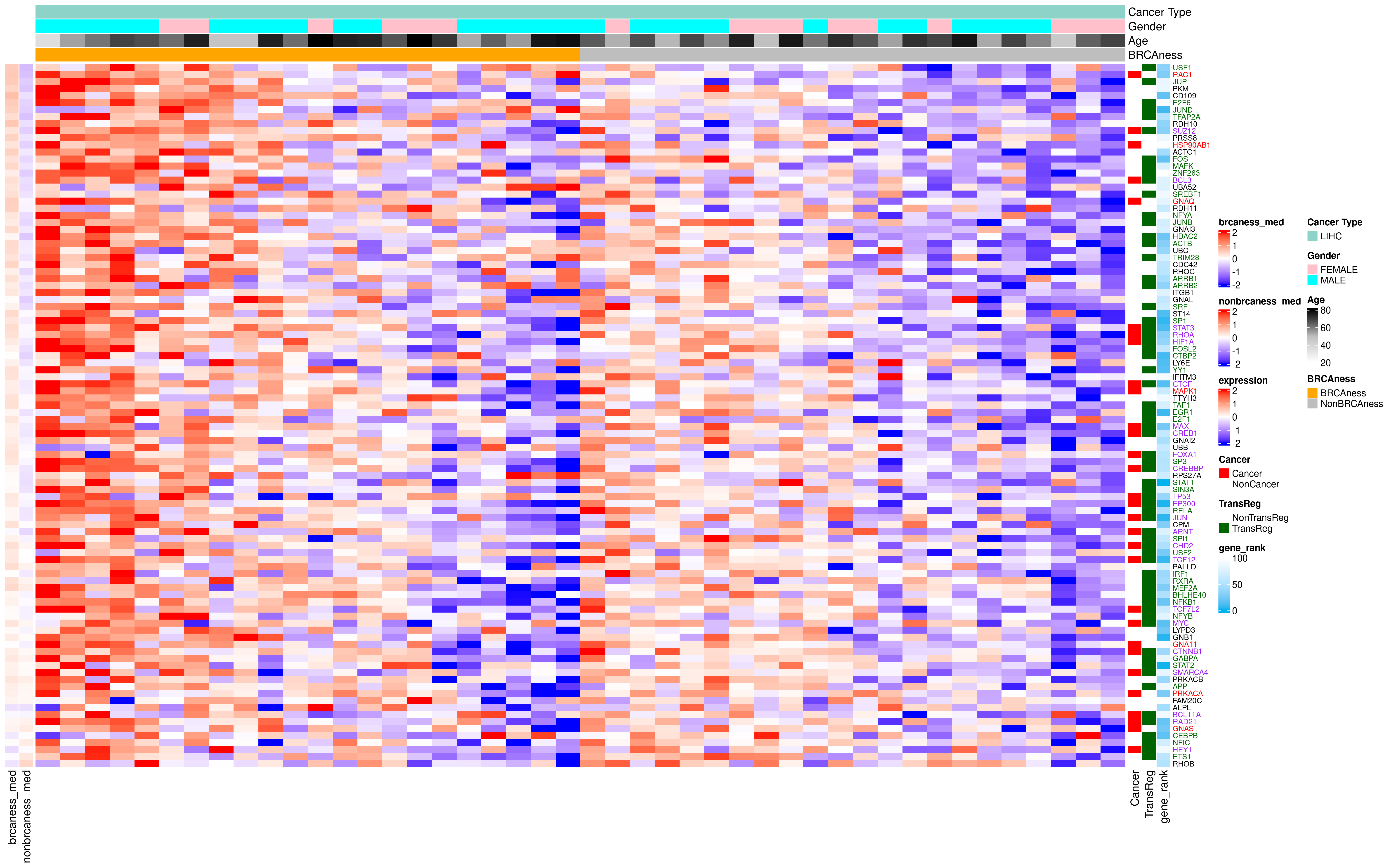

### LIHC_Network_Plot.png

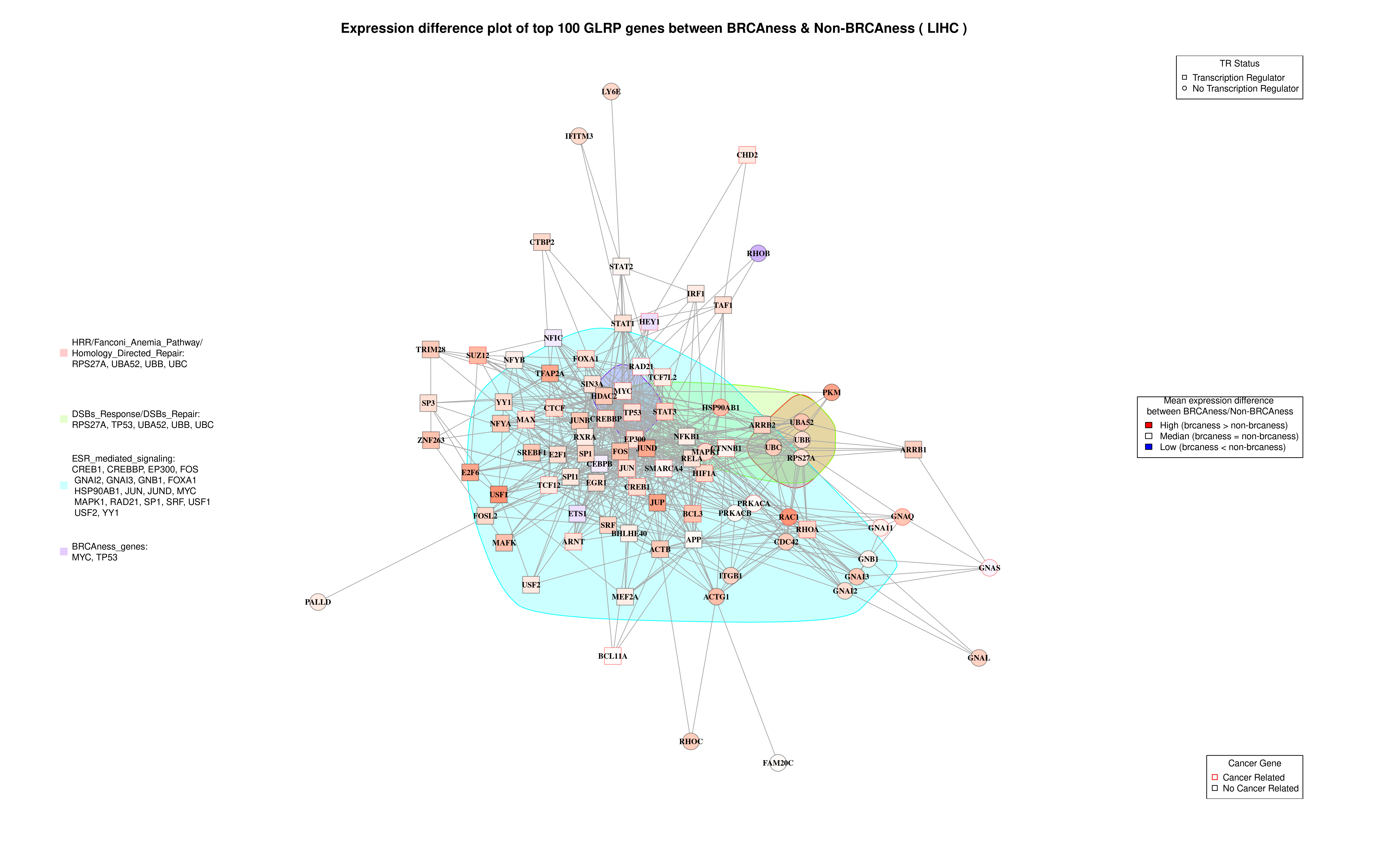

### LUAD_Heatmap.png

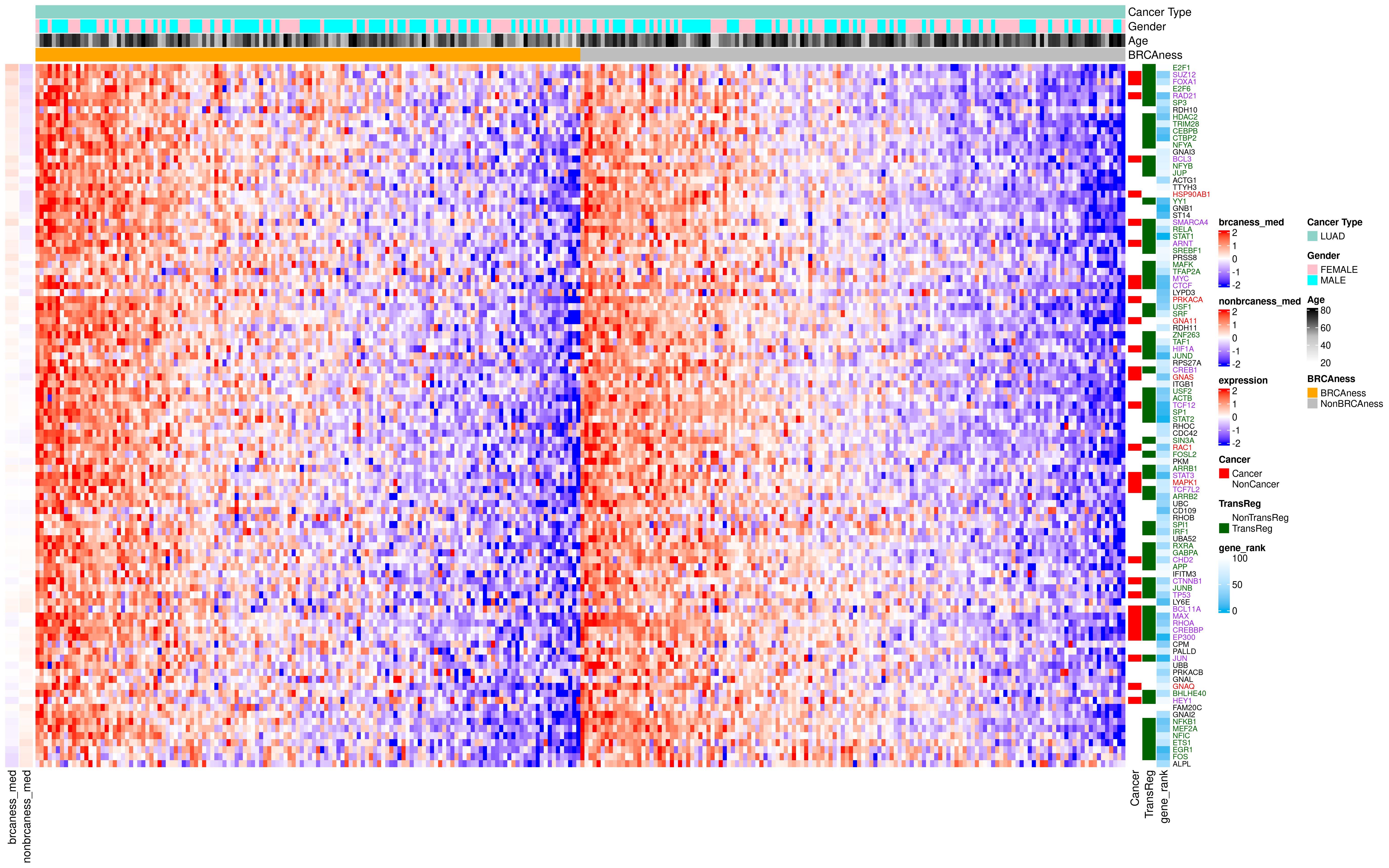

### LUAD_Network_Plot.png

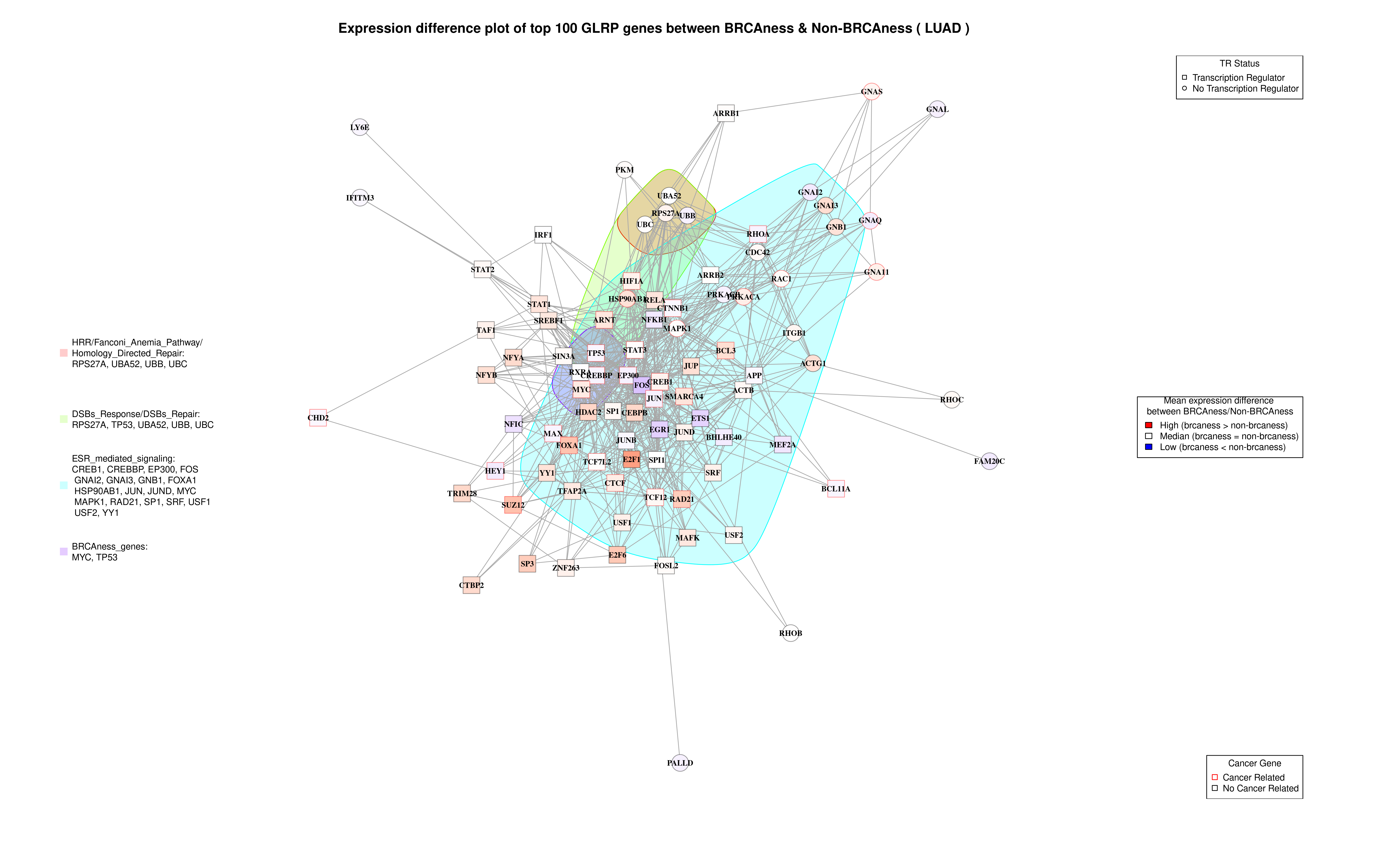

### LUSC_Heatmap.png

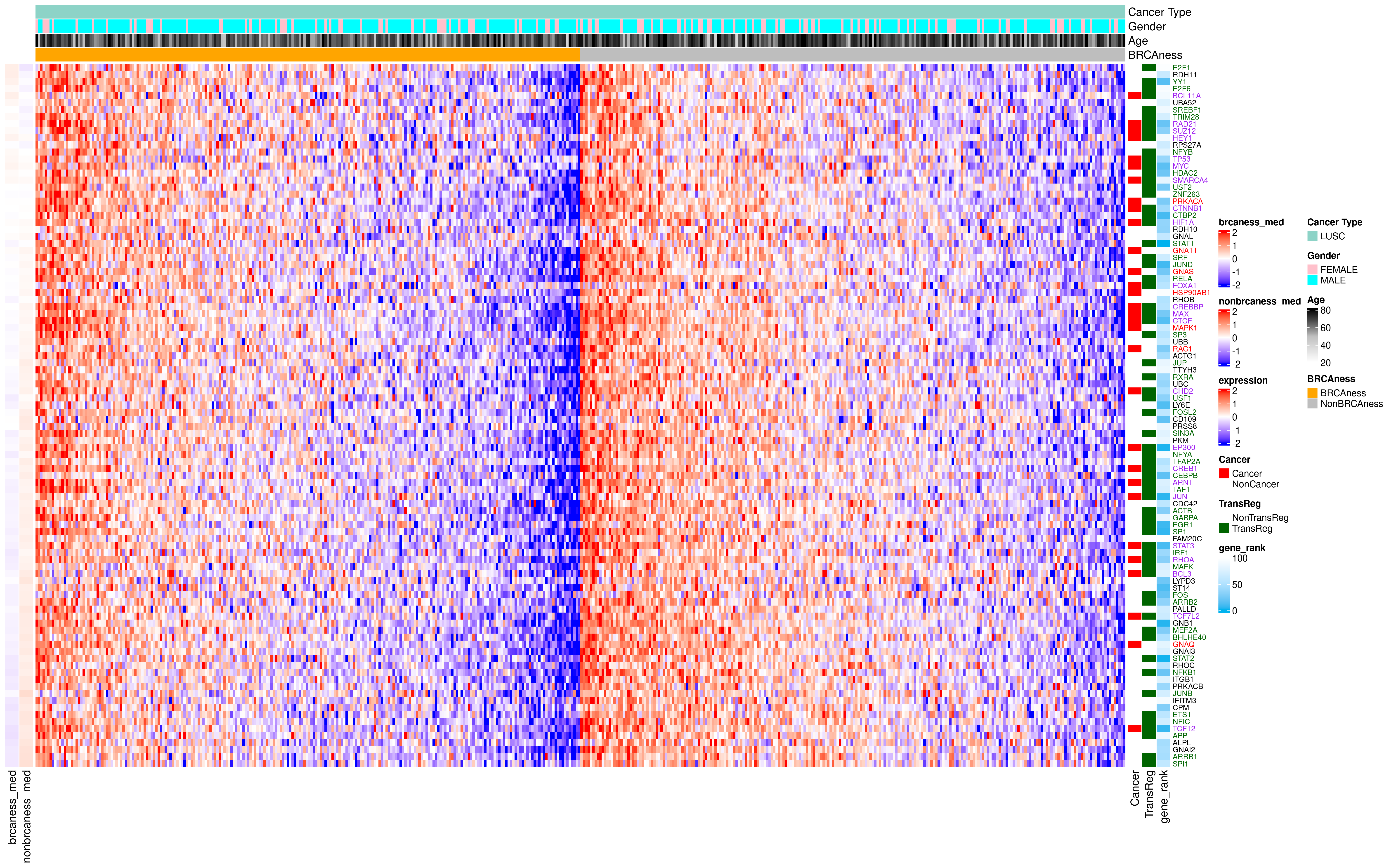

### LUSC_Network_Plot.png

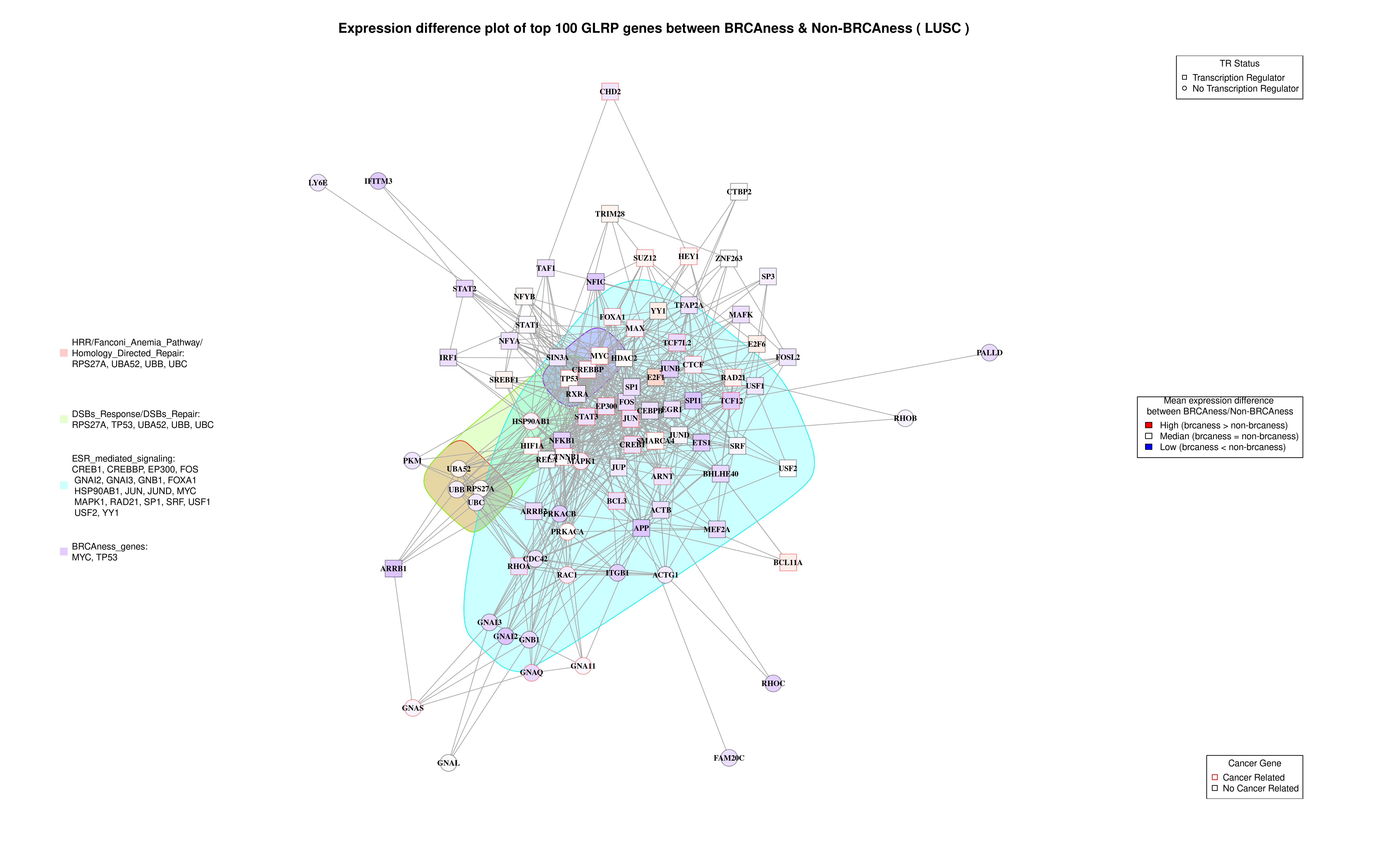

### OV_Heatmap.png

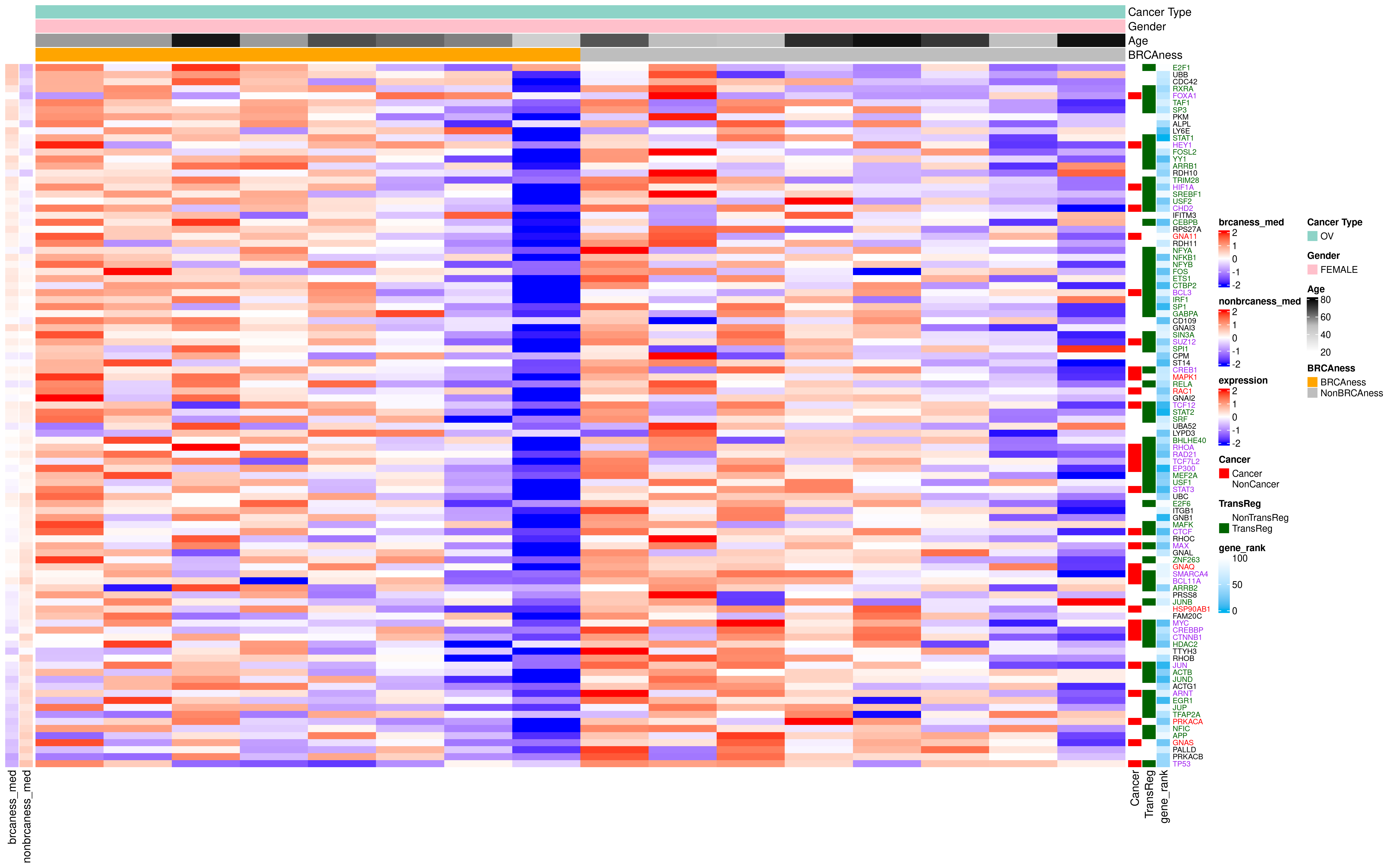

### OV_Network_Plot.png

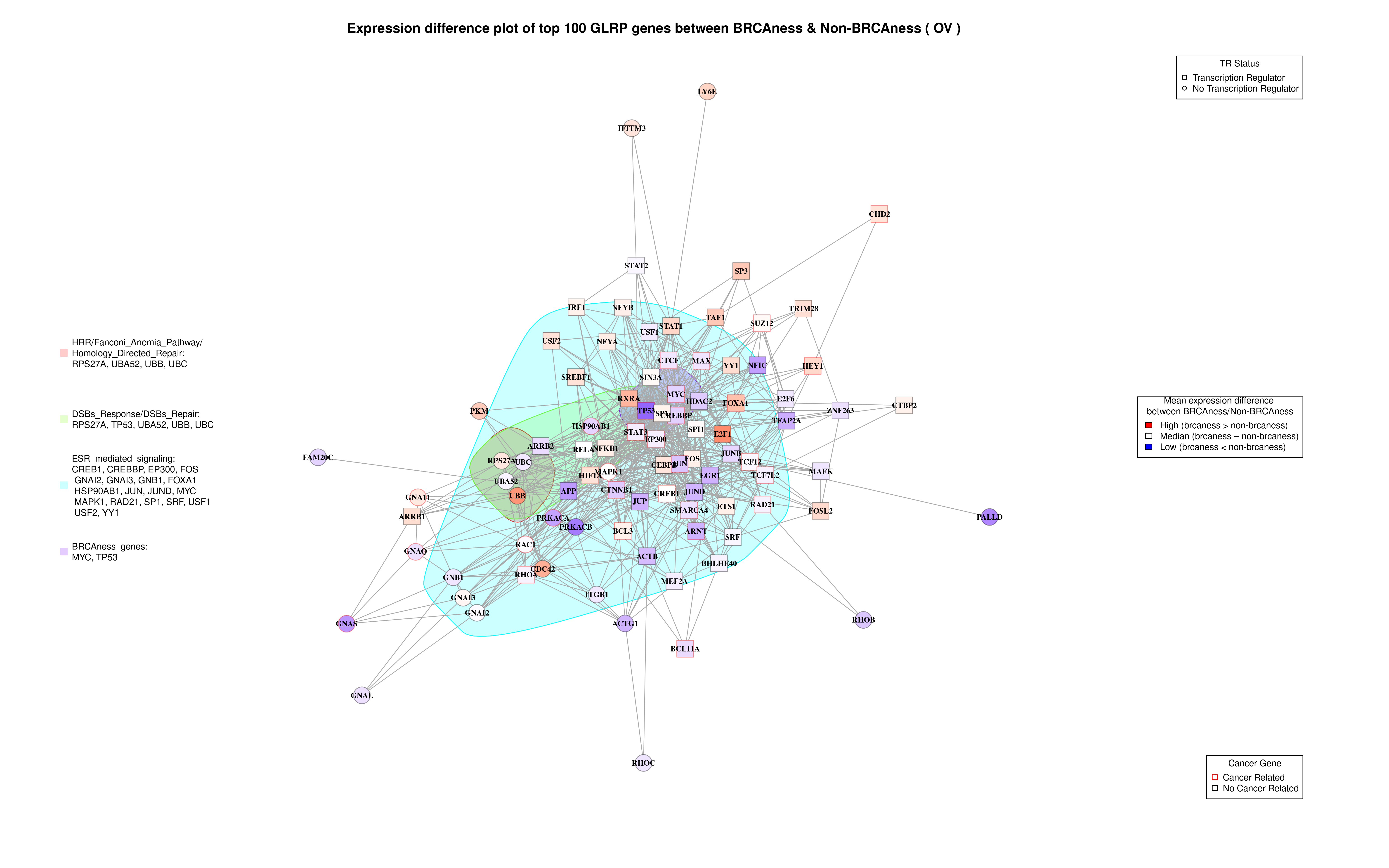

### PAAD_Heatmap.png

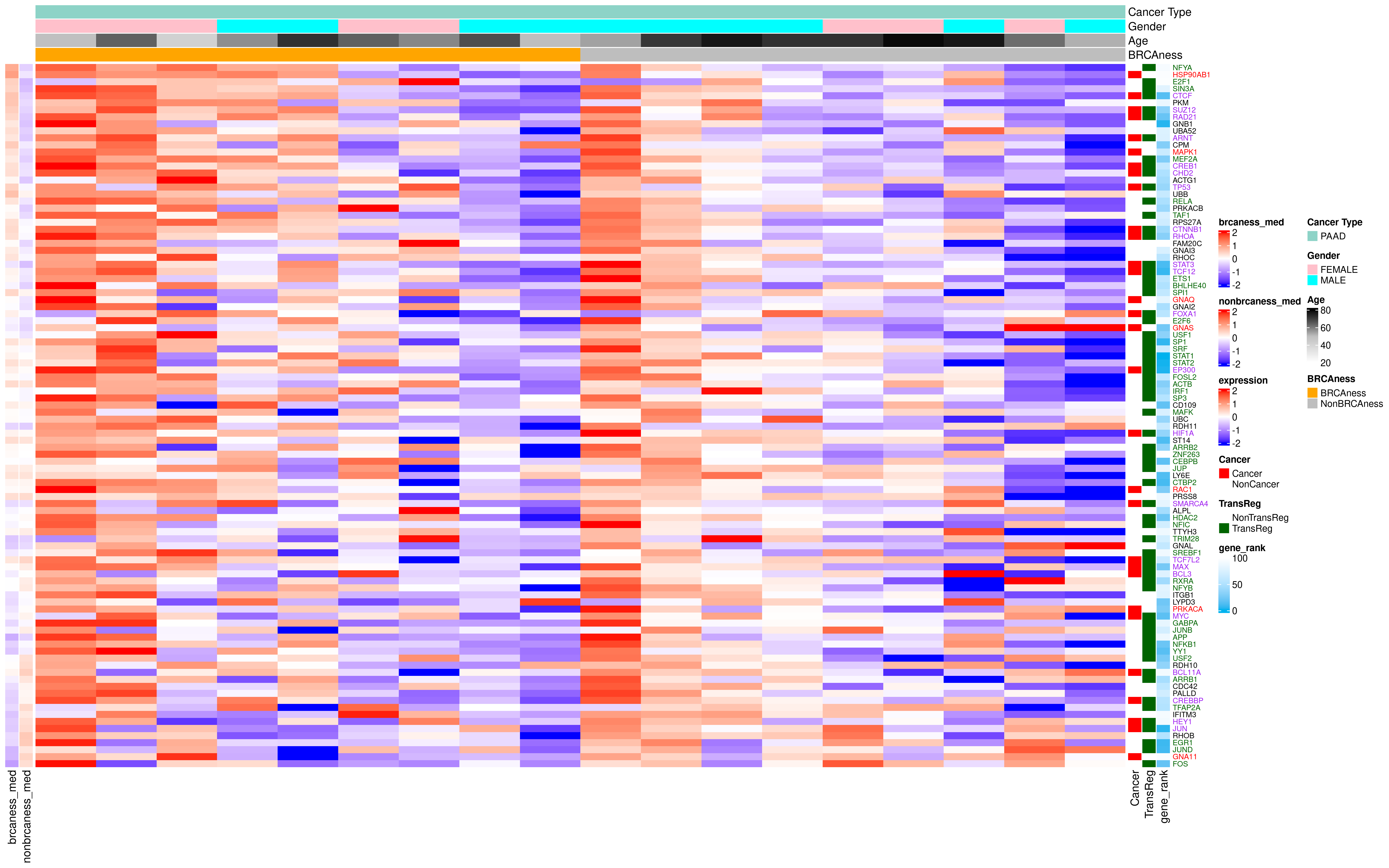

### PAAD_Network_Plot.png

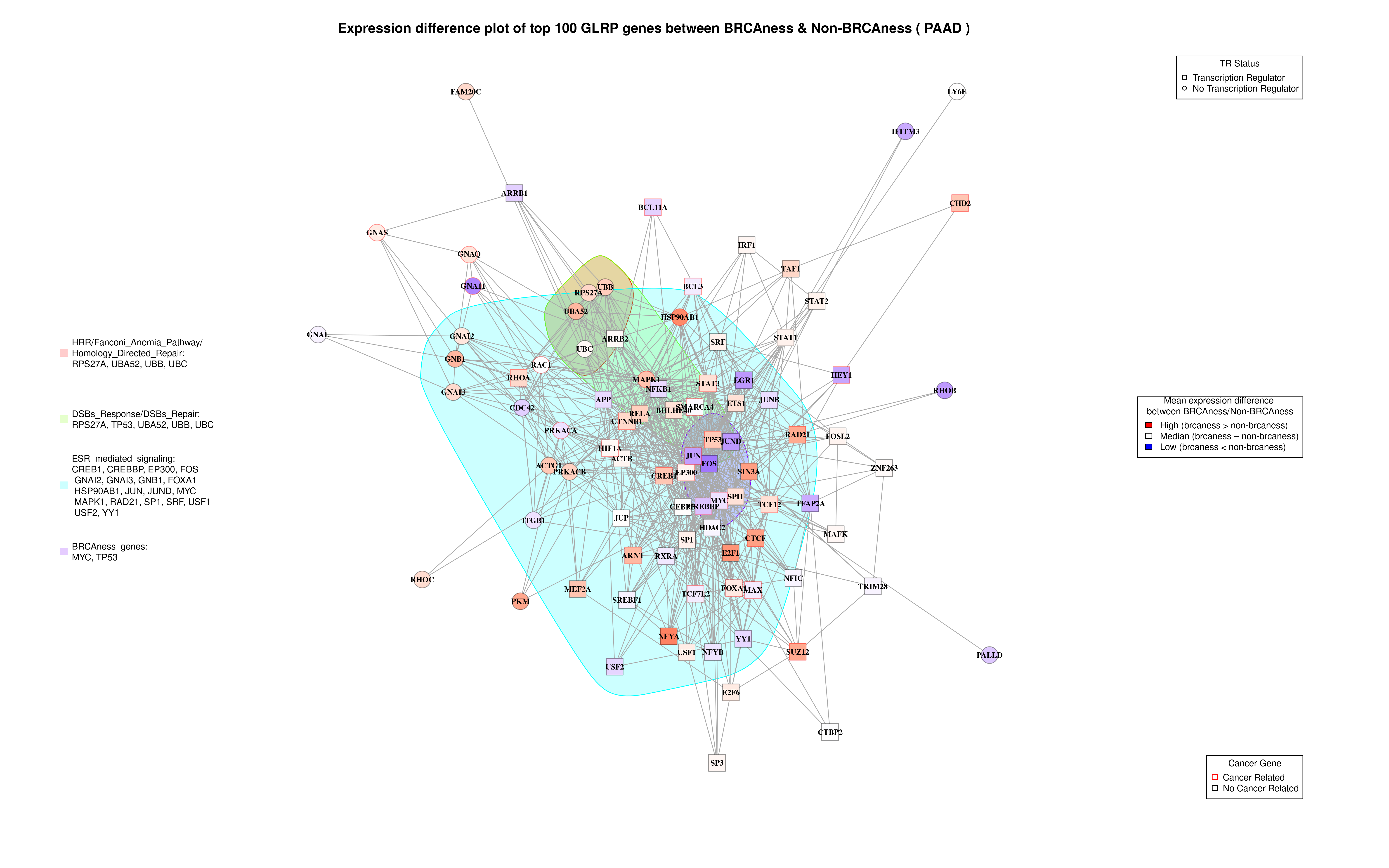

### PRAD_Heatmap.png

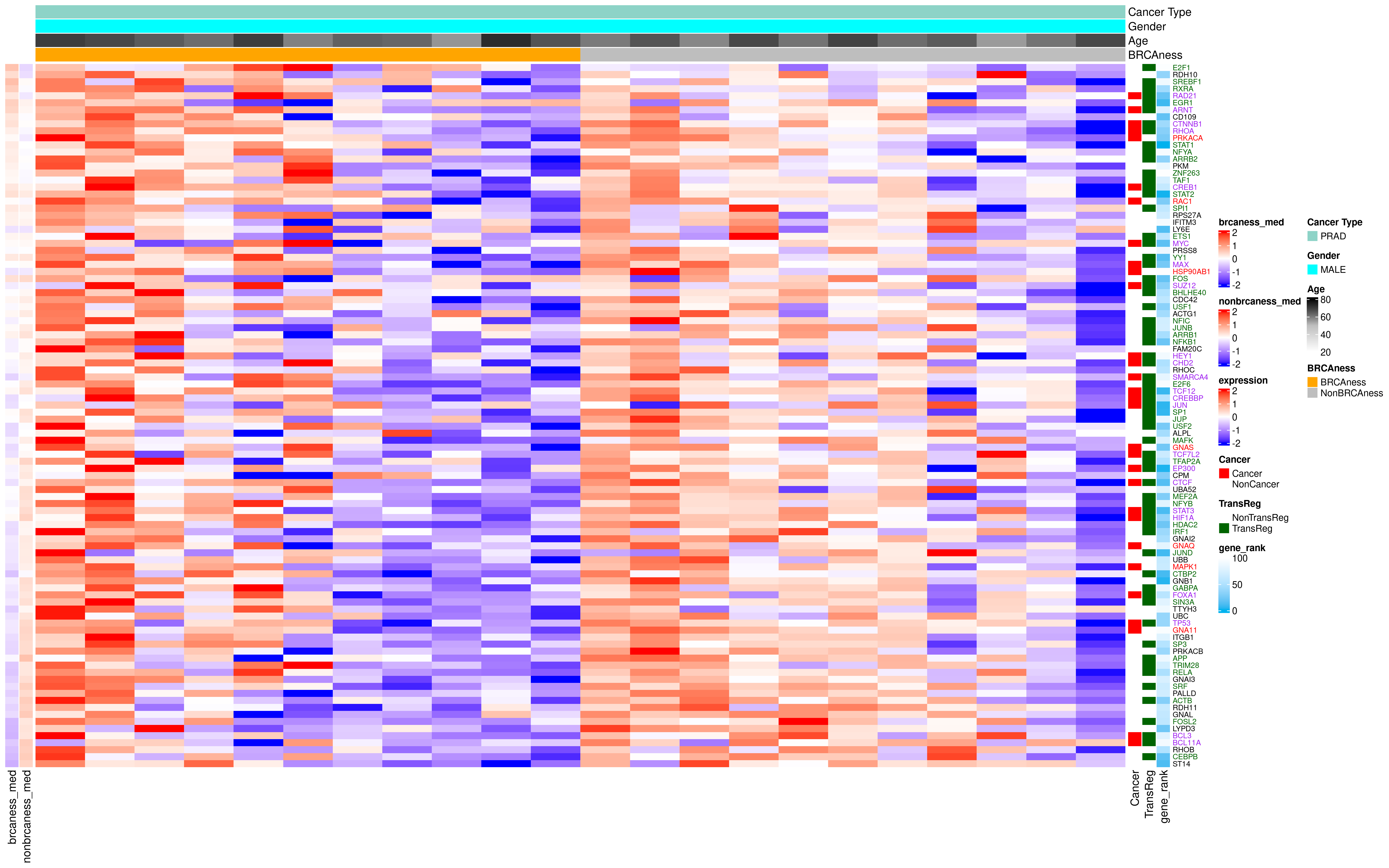

### PRAD_Network_Plot.png

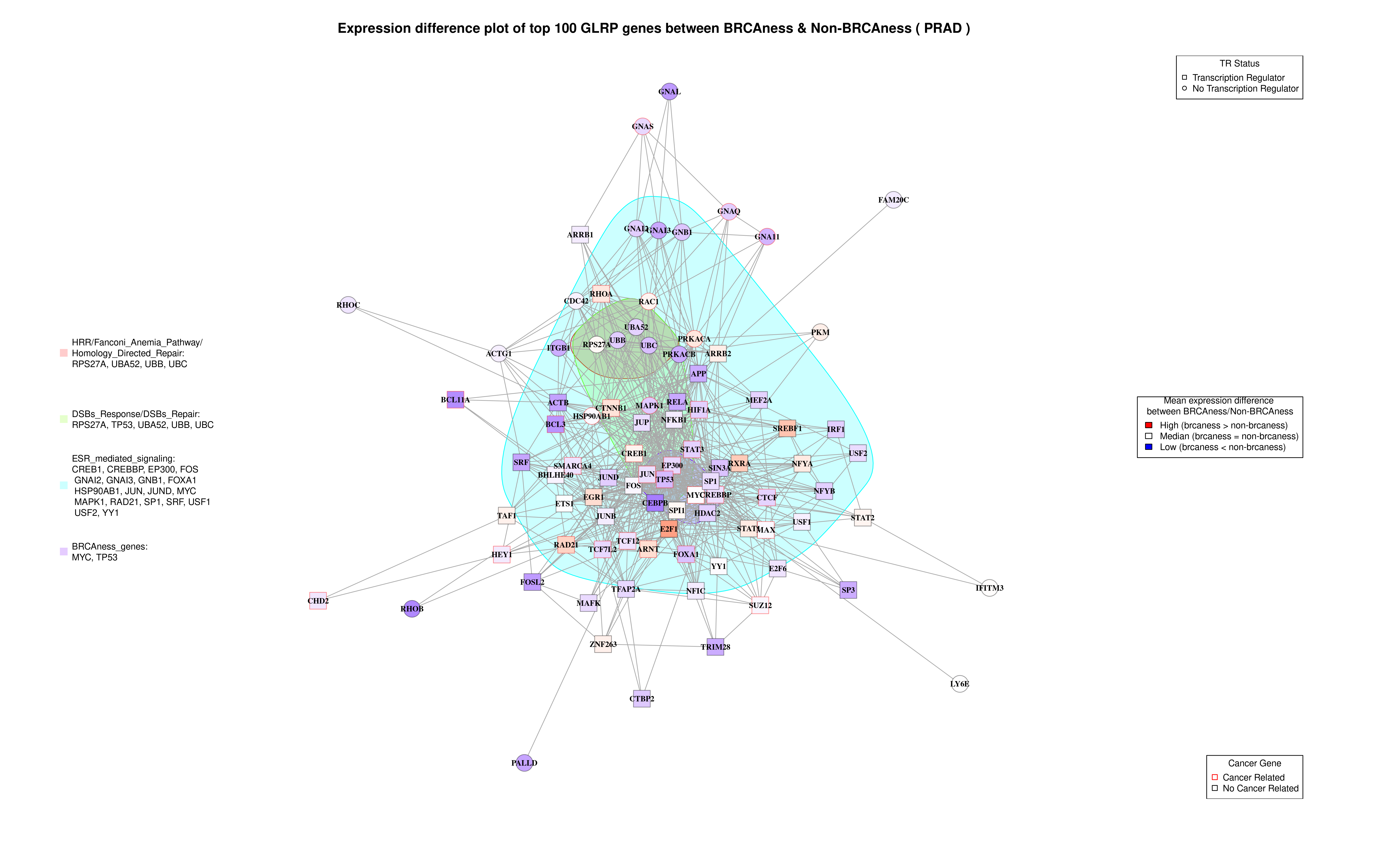

### READ_Heatmap.png

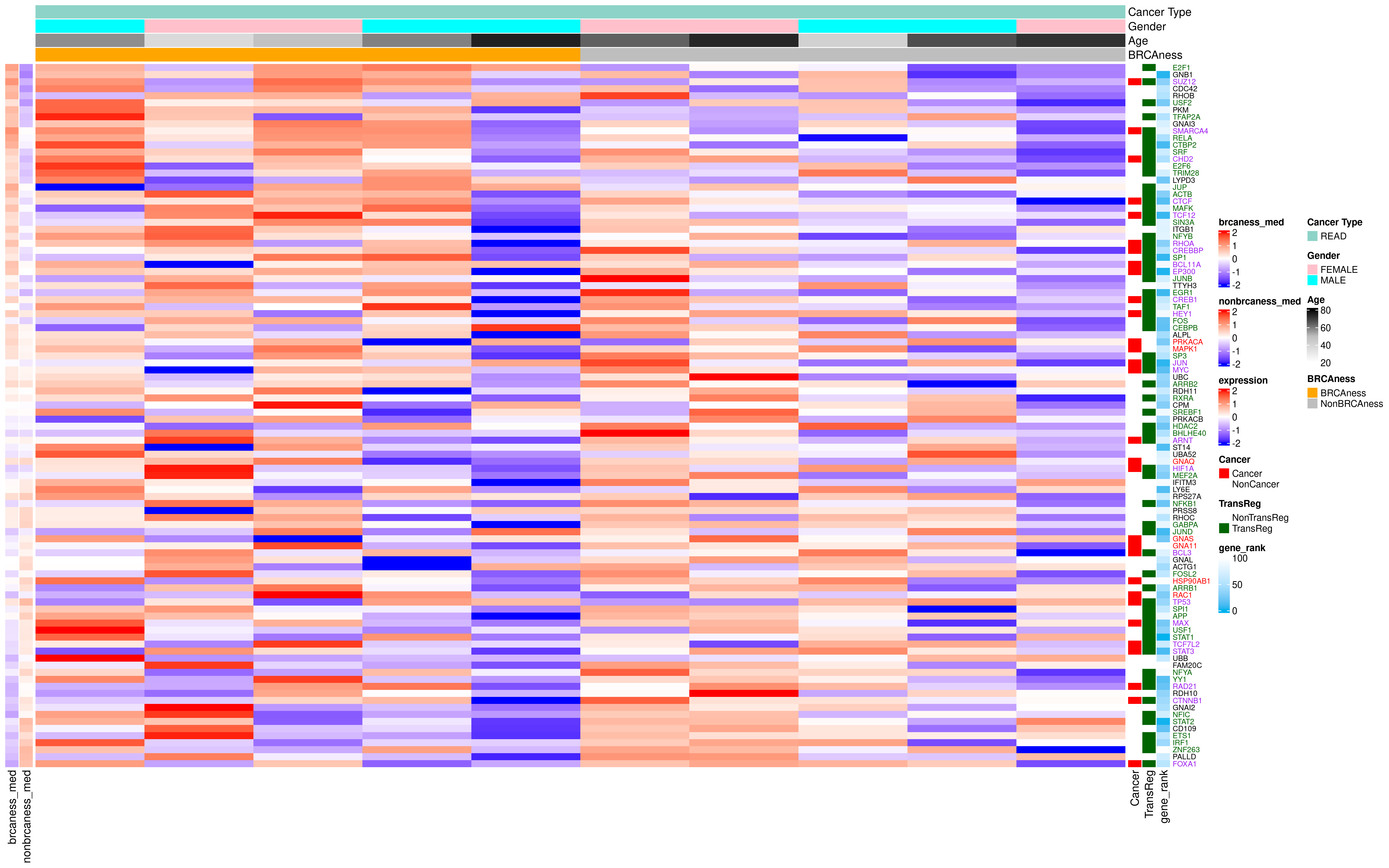

### READ_Network_Plot.png

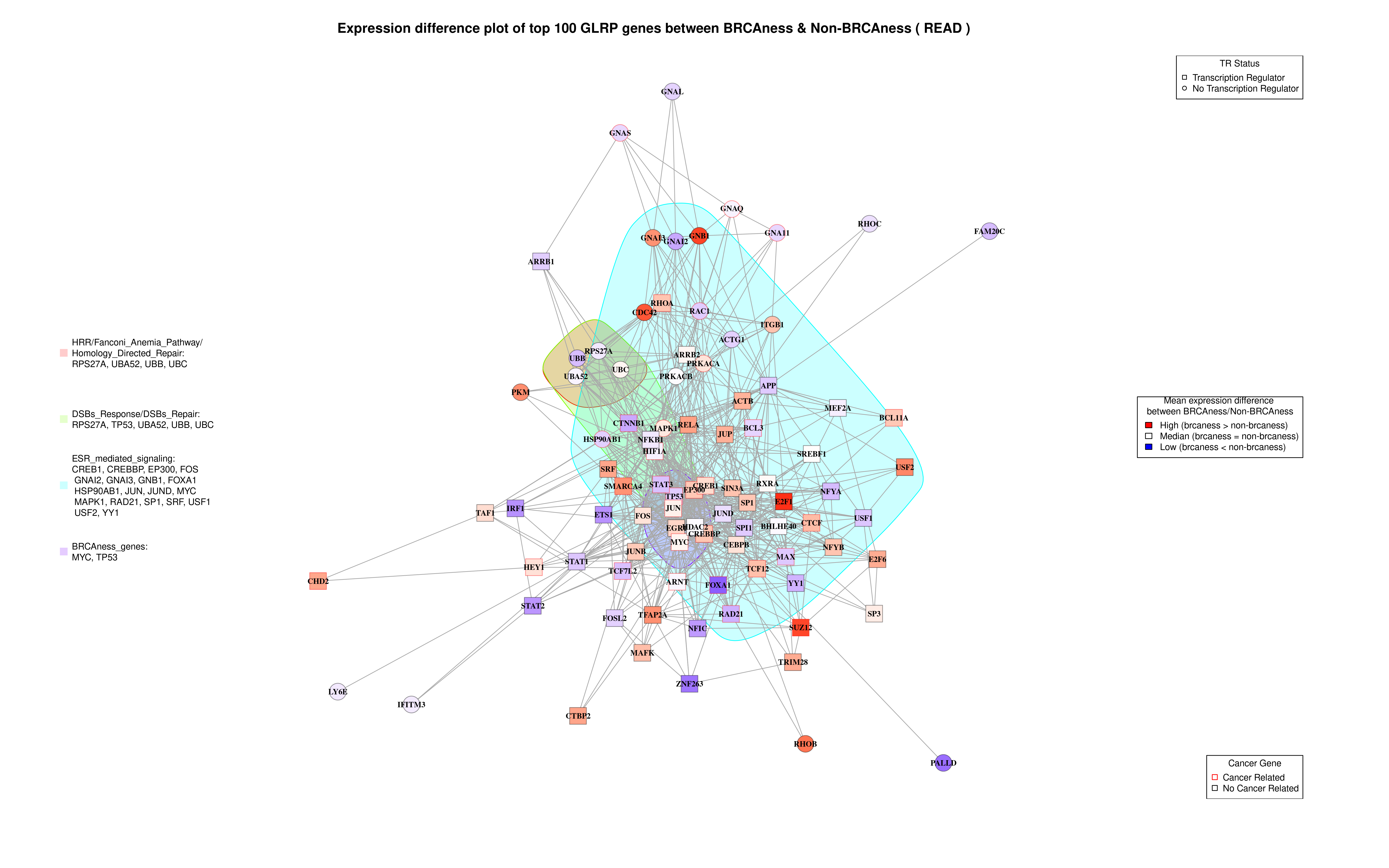

### SARC_Heatmap.png

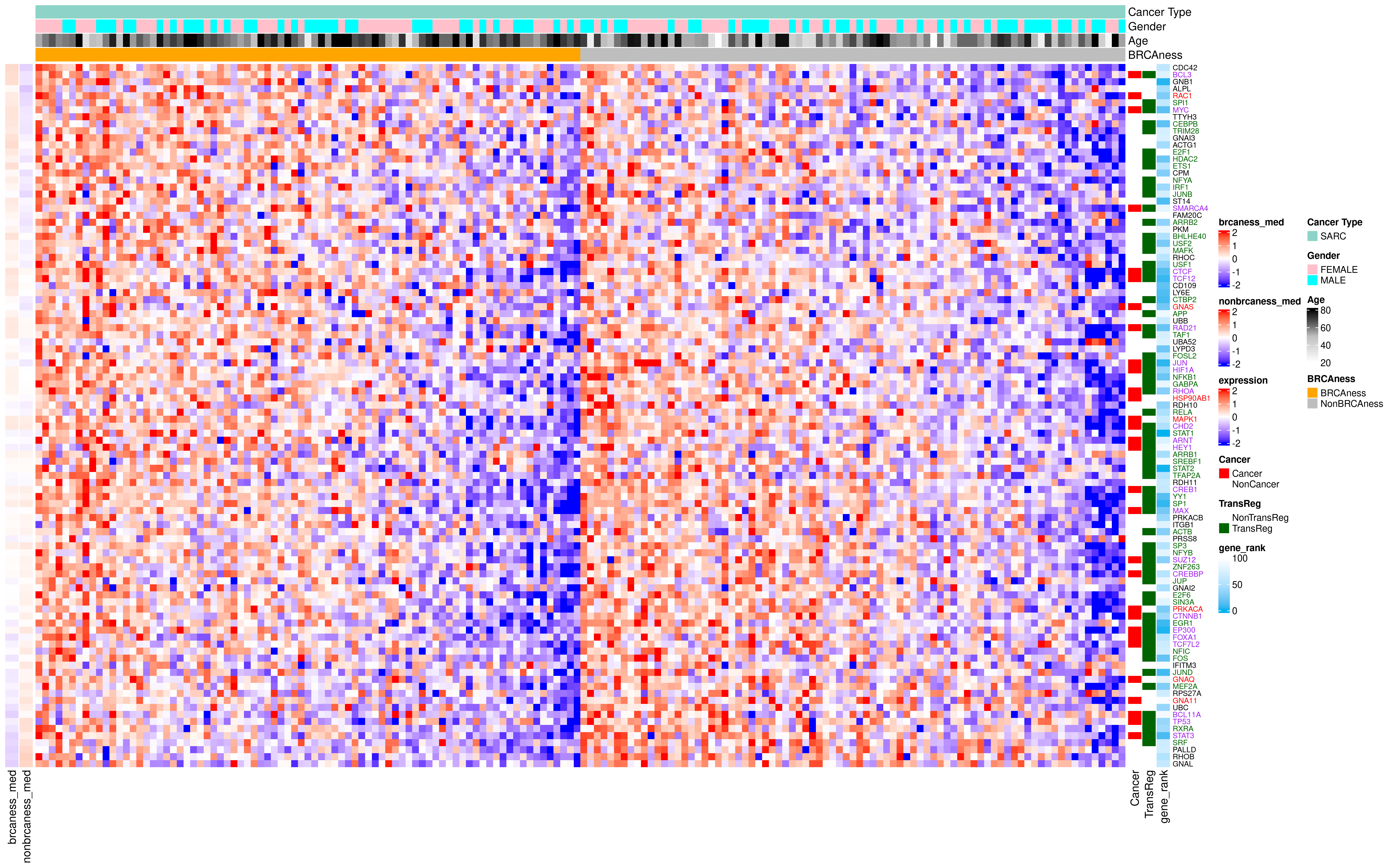

### SARC_Network_Plot.png

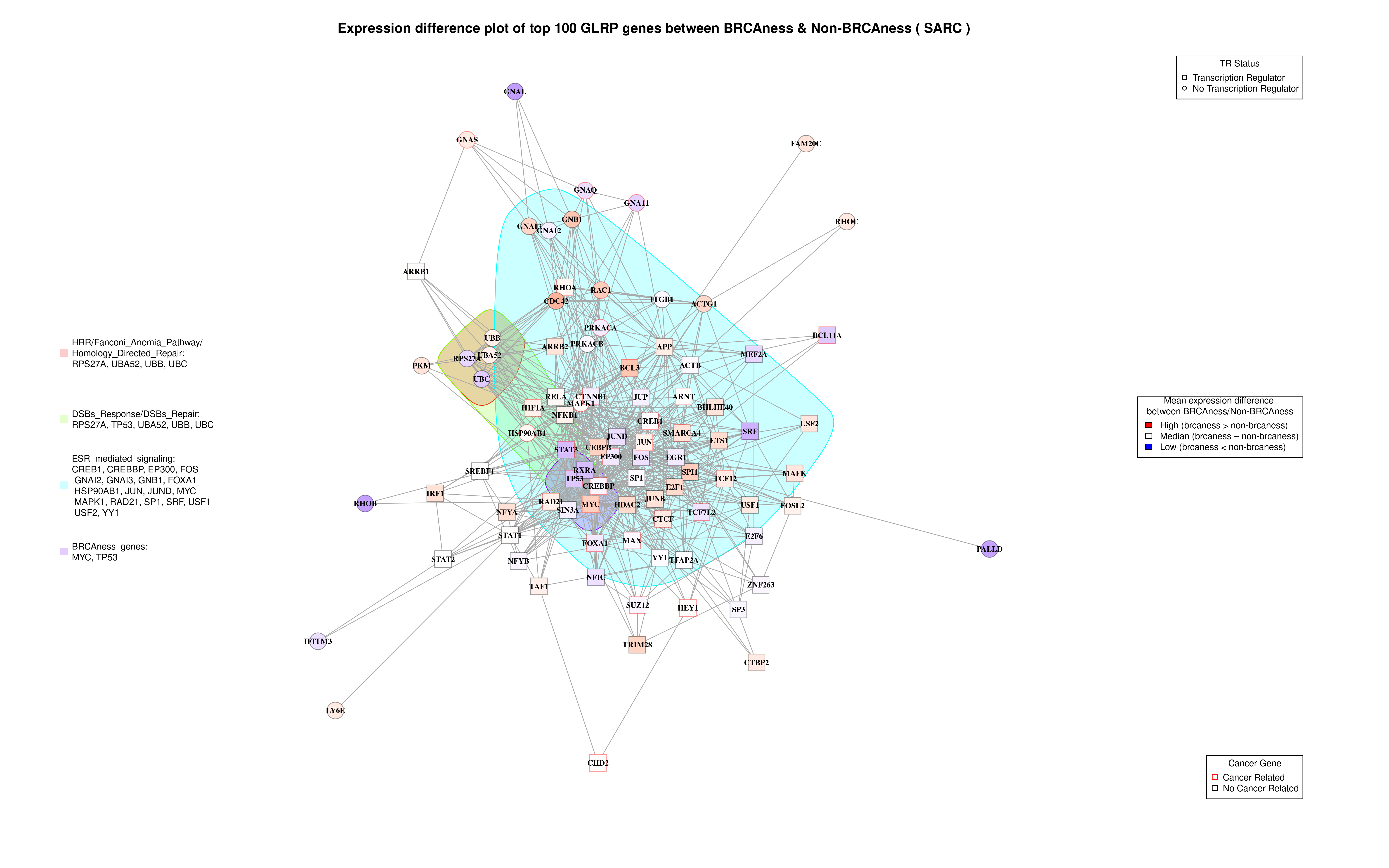

### SKCM_Heatmap.png

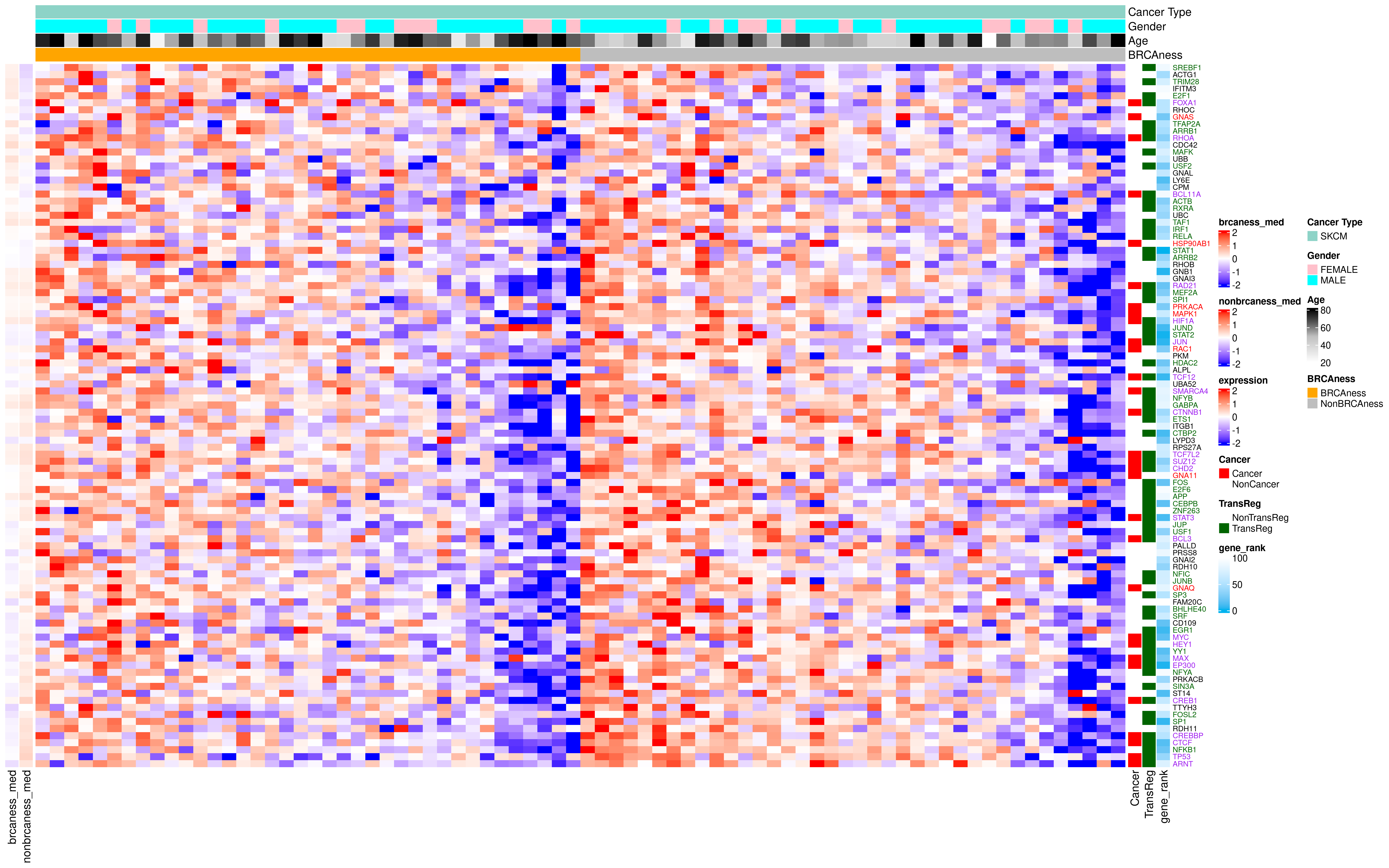

### SKCM_Network_Plot.png

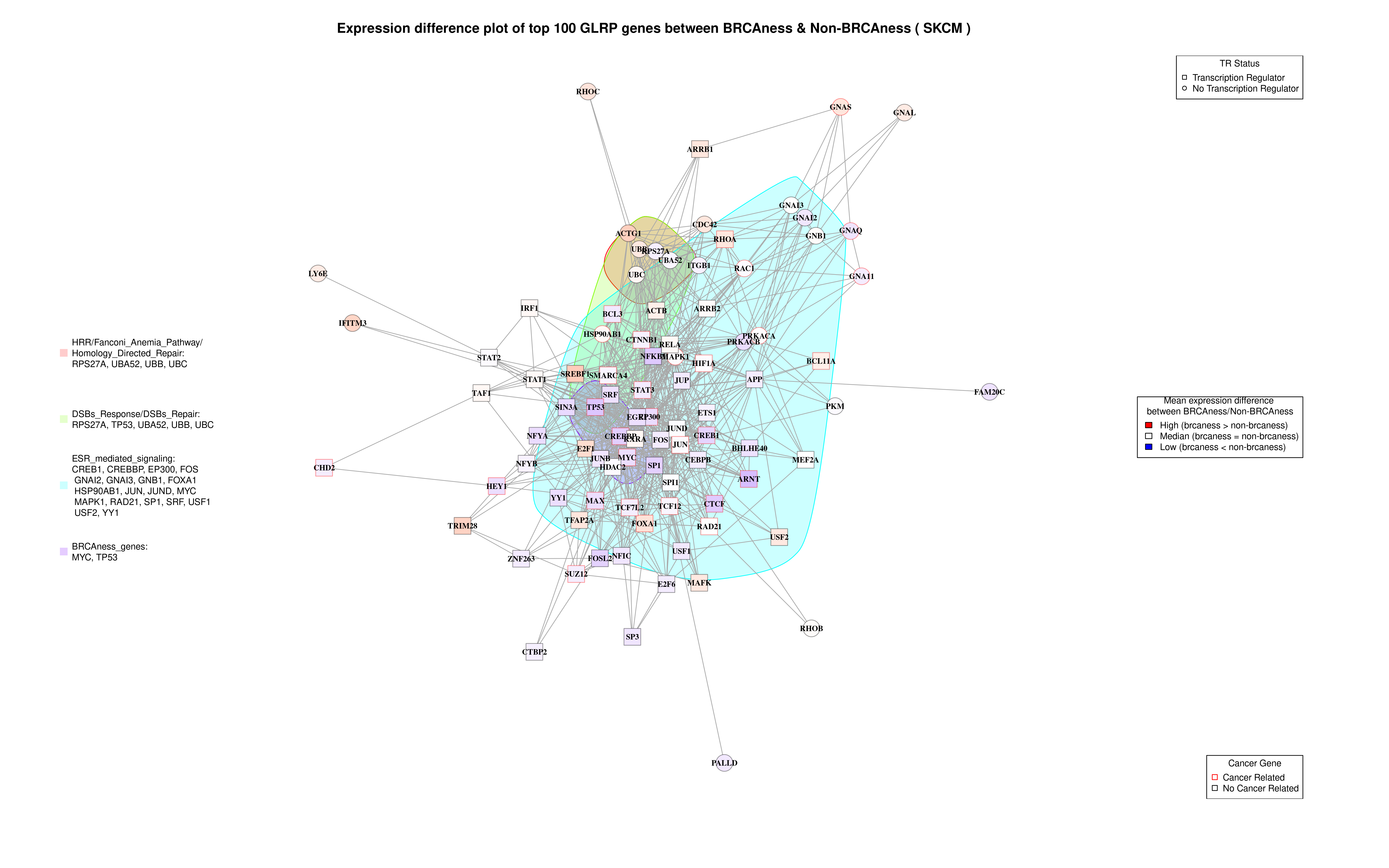

### STAD_Heatmap.png

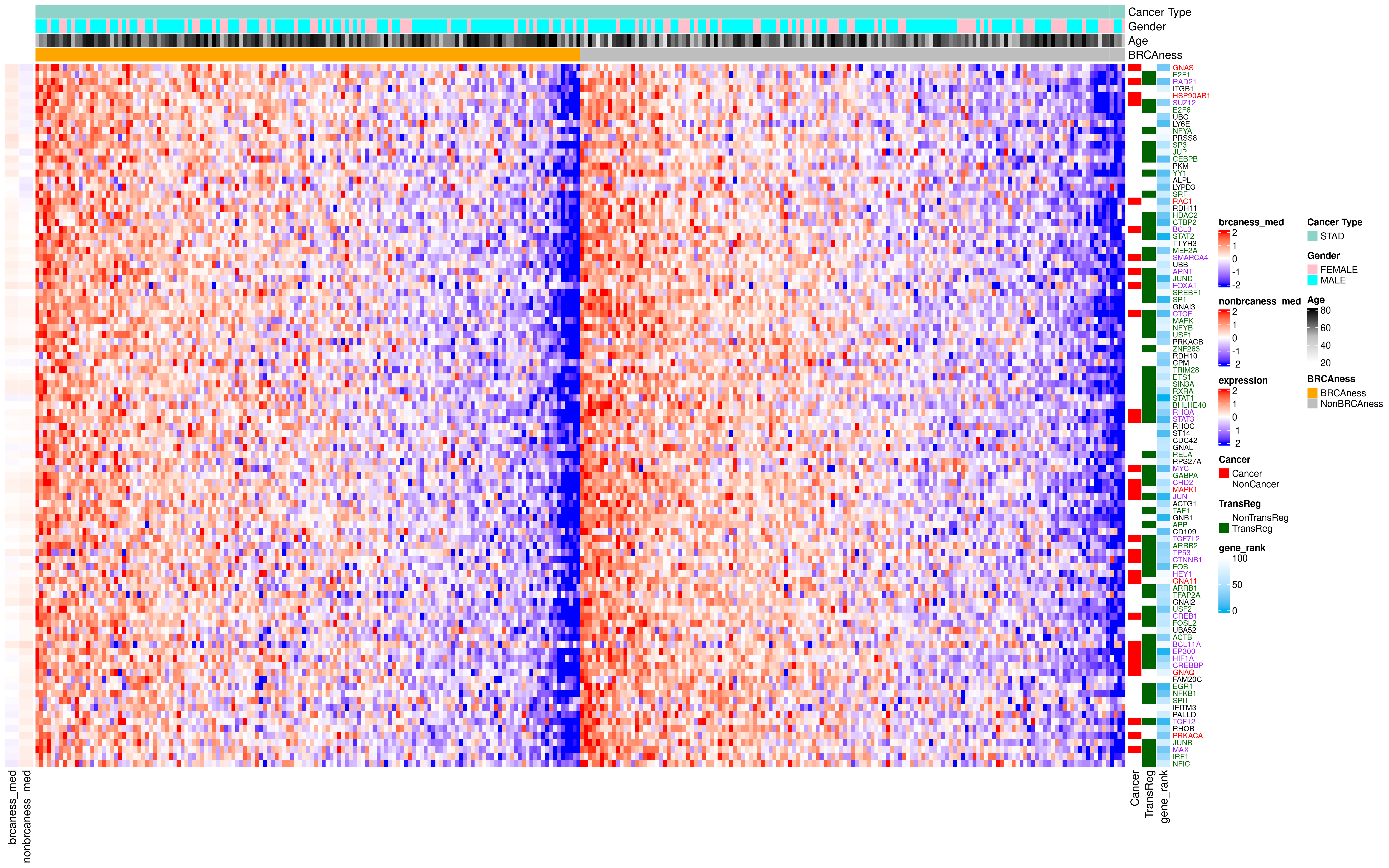

### STAD_Network_Plot.png

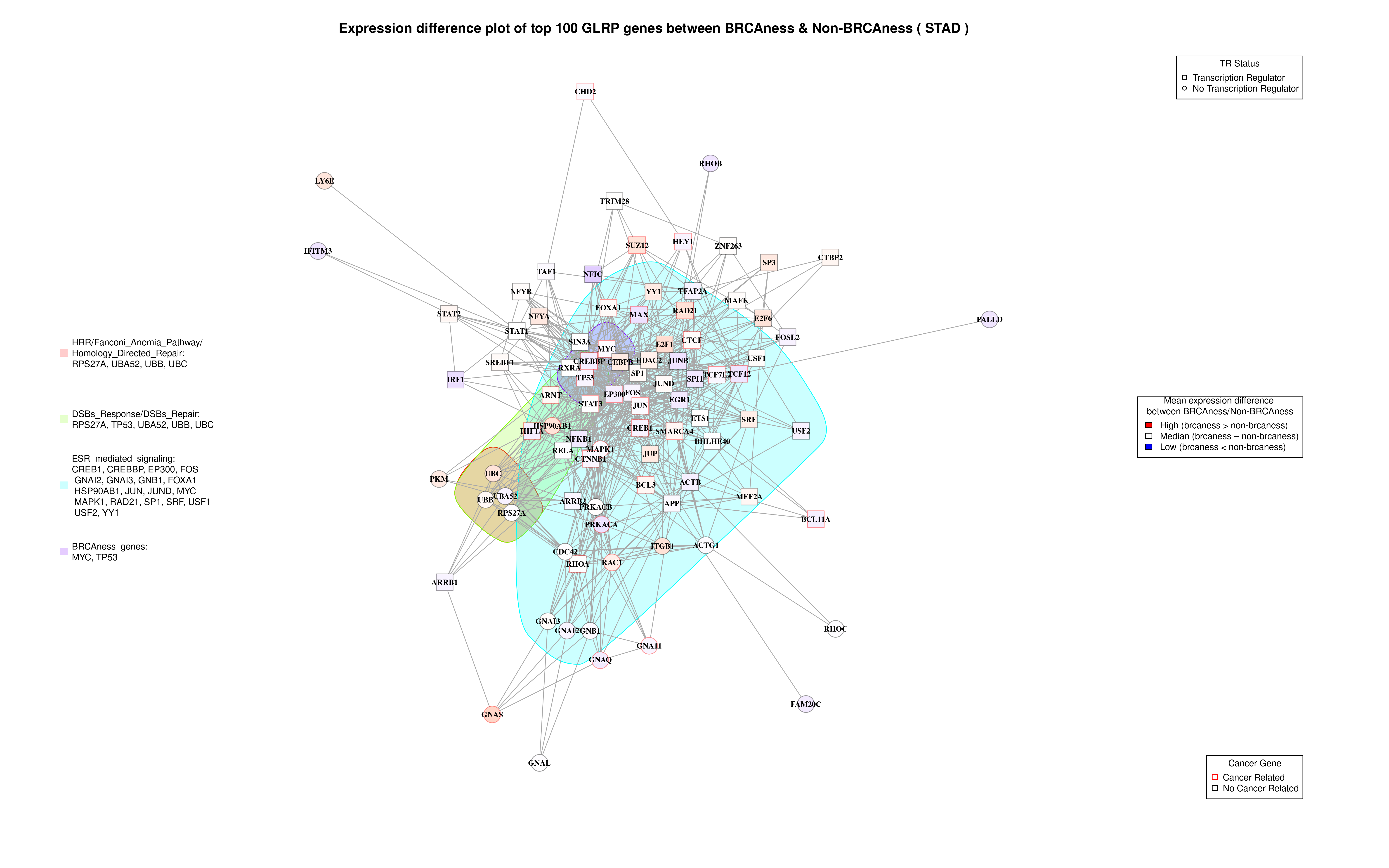

### UCEC_Heatmap.png

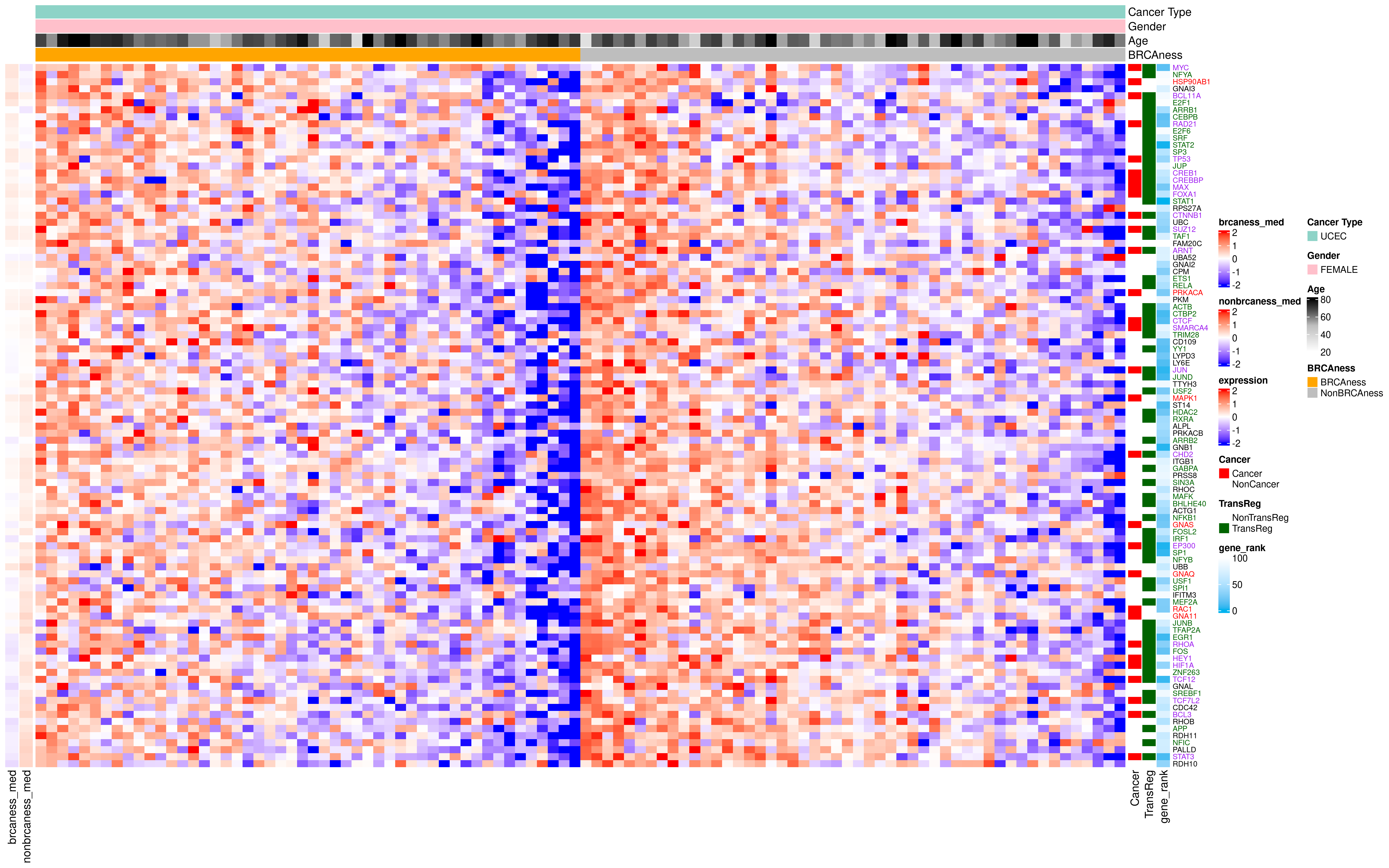

### UCEC_Network_Plot.png

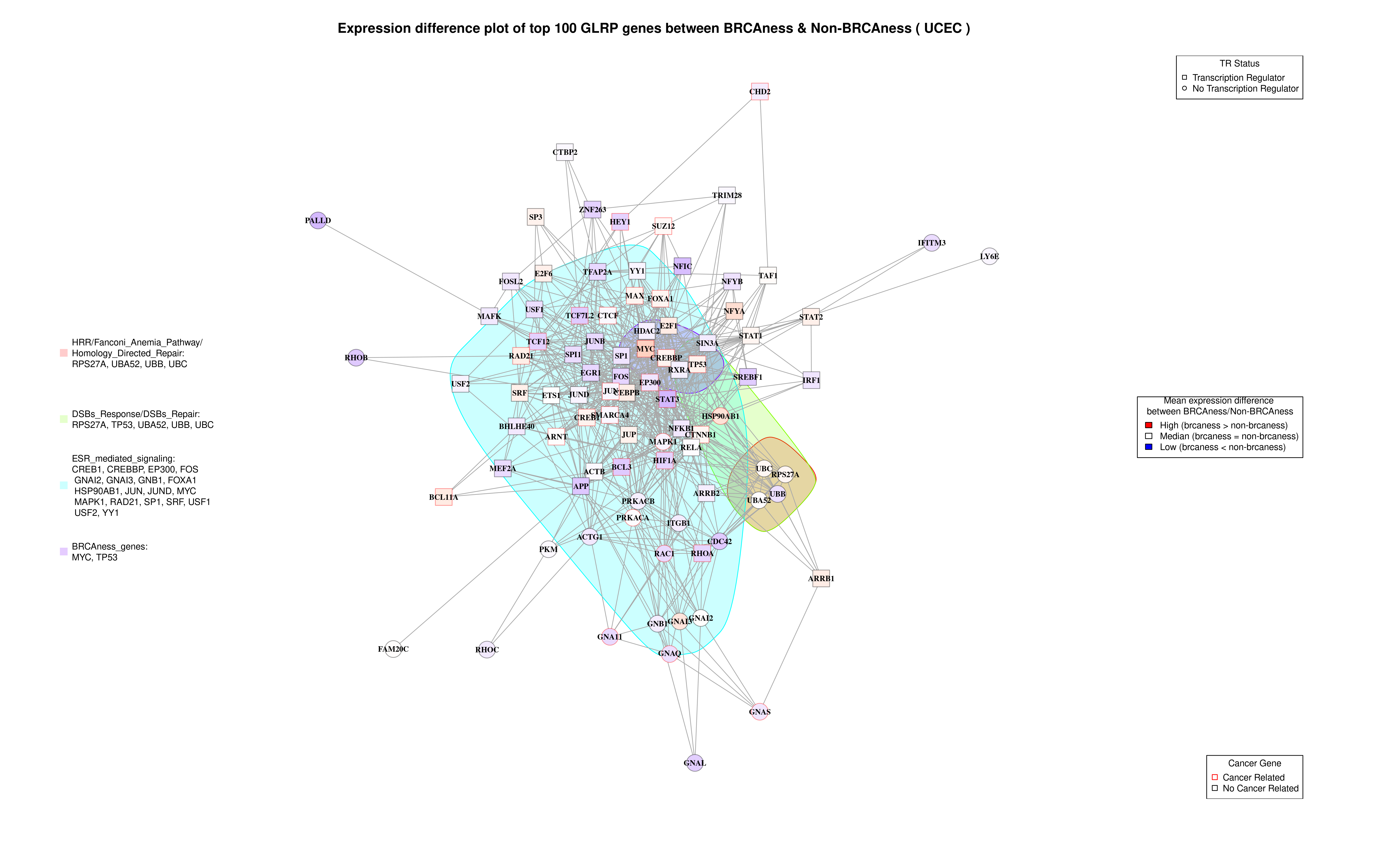

### UCS_Heatmap.png

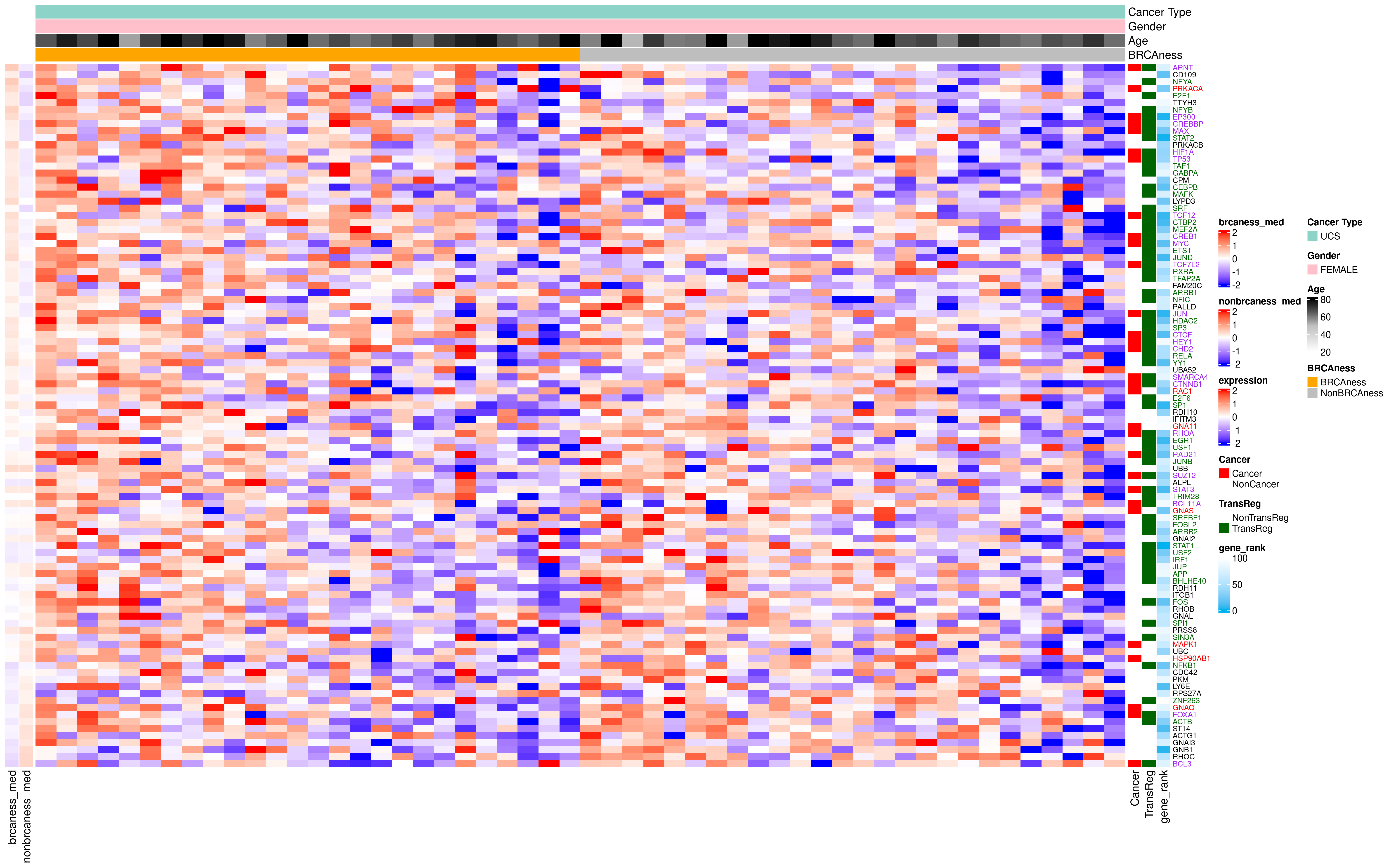

### UCS_Network_Plot.png

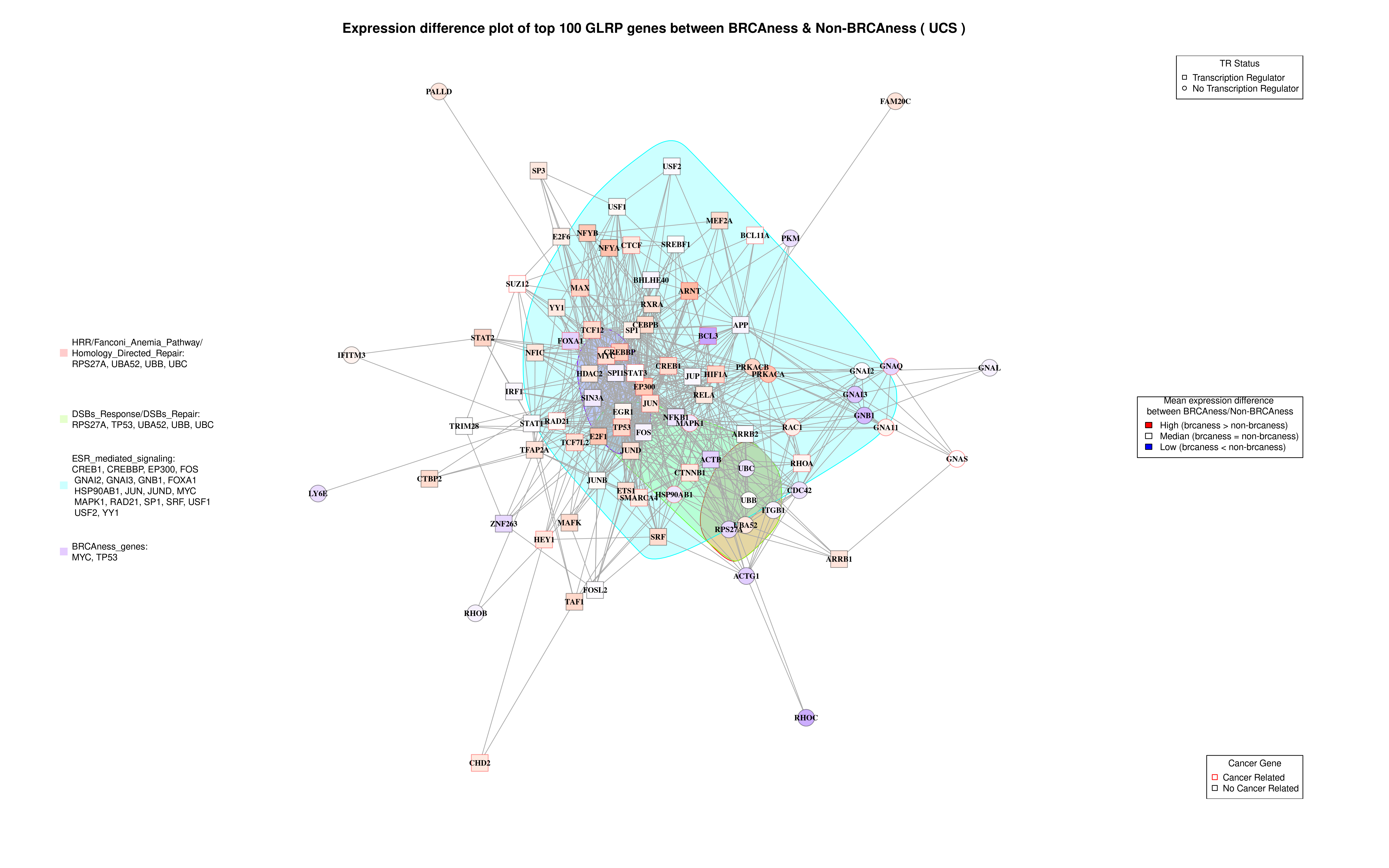
